# Talin-drug interaction reveals a key molecular determinant for biphasic mechanical effect: studied under single-molecule resolution

**DOI:** 10.1101/2022.04.04.486950

**Authors:** Soham Chakraborty, Madhu Bhatt, Debojyoti Chowdhury, Deep Chaudhuri, Shubhasis Haldar

**Author notes:** contributed equally to this work (Soham Chakraborty, Madhu Bhatt, Debojyoti Chowdhury).

## Abstract

Talin as an adhesion protein, exhibits a strong force-dependent structure-function dynamics. Being a mechanosensitive focal adhesion (FA) protein, talin might interact to several FA targeting drugs; however, the molecular mechanism of talin-drug interactions remains elusive. Here we combined magnetic tweezers and molecular dynamics (MD) simulation to explore mechanical stability of talin with three drugs based on their talin specificity. Interestingly, our study revealed that talin displays a bimodal force distribution with a low and high unfolding force population. We observed that talin nonspecific drugs (tamoxifen and letrozole) display biphasic effect: increase talin mechanical stability upto optimum concentration, followed by a decrease in stability with further concentration increase. By contrast, talin-specific cyanidin 3-O-glucoside promotes a steady increase to talin mechanical stability with its concentration. We reconciled our observation from the simulation study: tamoxifen enters into talin hydrophobic core, eventually destabilizing the protein; whereas cyanidin 3-O-glucoside stabilizes the protein core by maintaining the inter-helix distance. Finally, we observed a strong correlation among hydrophobicity and cavity analysis, illustrating a detailed mechanistic analysis of drug effect on the mechanosensitive protein. Overall this study presents a novel perspective for drug designing against mechanosensitive proteins and studying off-target effects of already known drugs.

## Introduction

Adhesion proteins, both at cell-matrix and cell-cell interfaces, control diverse cellular processes ranging from cell migration to its structural maintenance through a comprehensive force governed mechanism called mechanotransduction^1–4^. Structure-function dynamics of these proteins are strongly dependent on mechanical stability, i.e., their capability to withstand force, generated during cellular traction^3,4^. Since these adhesion complexes are junctions of pathological cell migration, force-responses of these individual proteins are in prime focus for therapeutic applications in different diseases. Studies have reported integrins, microtubule and actin are currently used as potential drug targets. Similarly, focal adhesion kinase (FAK) degraders have been successfully developed through different methodologies including PROTAC technology^5,6^. At molecular level, these small molecule drugs could modulate the structural stability of these proteins, which in turn could affect their mechanical response and protein interactions.

Mechanosensitive adhesion proteins are structurally predominated by α-helix folds, which acts as a critical determinant of their forced unfolding and mechanical stability: myosin II, which mediates FA maturation under force, are inhibited by N-benzyl-p-toluene sulphonamide within micromolar concentration range. Similarly, F-actin has been observed to interact with various drugs including flutamide, carvedilol and mitoxantrone^7–9^. Recently, cytochalasin-D and vinblastine as cytoskeletal drugs regulate off-target folding stability of other protein in cells^10^. Furthermore, tamoxifen treatment regulates cell adhesion via phosphorylating FAK and p130Cas in focal adhesion^11,12^. Catenin proteins, containing a 4-helix conformation, exhibit mechano-regulatory functions and are plausible drug targets in pathological cell migration.^13–15^ Similarly, talin is a key α-helical FA protein, which exhibit strong force-dependent folding dynamics and concurrent interactions with other proteins such as FAK, integrin, and paxillin^16–18^. These talin-centred mechanical linkages act as crossroads of pathological cell migration and proliferation and could act as a plausible drug target. Therefore, adhesion targeting drugs could have pronounced effect on talin mechanical stability, which could further be tuned by substrate specificity of drugs.

Here we combine single-molecule magnetic tweezers and molecular dynamics (MD) simulation to demonstrate the drug effect on a mechanically stable talin R3-IVVI domain, which is predominantly α-helical and have been used as a model substrate in several force spectroscopic studies^18–21^. Our force spectroscopic results revealed that talin unfolding distinctly exhibits a bi-modal force histogram with rare unfolding events at higher force range. At different concentrations, these drug molecules could modulate the relative population of both the low and high force unfolding events. To investigate their effect on the mechanical stability, we selected three drugs on the basis of the talin specificity: tamoxifen, letrozole, and cyanidin 3-O-glucoside, which have been reported to affect cell adhesion in different pathological conditions. Tamoxifen and letrozole have not been reported to interact with talin, while cyanidin 3-O-glucoside binds to talin F3 domain. Interestingly, we observed tamoxifen and letrozole display biphasic effect, promoting higher mechanical stability upto optimal range; followed by decreasing the stability with the further increase in the concentration. However, this effect was not experienced with talin-specific drug cyanidin 3-O-glucoside, where we found a steady increase of the high force population and thereby, higher mechanical stability with the concentration. To gain underlying molecular mechanism of this key finding, we further performed MD simulation with these two drugs: tamoxifen molecules enter the hydrophobic core of the talin, eventually displacing their inter helix distance at high concentration, whereas cyanidin 3-O-glucoside maintains the inter-helix distance and radius of gyration, which stabilizes the talin domain. Finally, we are able to correlate the hydrophobicity, cavity analysis and the amount of penetrating drug molecules into the core to illustrate a comprehensive picture of drug effect on the mechanosensitive proteins. Overall, this detailed molecular insight into drug interaction with mechanosensitive protein could have significant implication for redesigning their mechanical stability as a plausible therapeutic target for adhesion targeting drugs.

## Results

### Distinct mechanical response of talin probed by magnetic tweezers

In our experimental set-up, the talin domain is flanked by N-terminal HaloTag for the covalent attachment on the glass surface, and a C-terminal AviTag for interacting to the paramagnetic bead through biotin-streptavidin chemistry. Force are applied by generating a magnetic field through permanent magnets (Fig. 1A). The unfolding force was measured by a force-increase scan with a loading rate of 3.1 pN/s. We observed the unfolding event as a sudden increase in the extension at a particular force. The unfolding steps are vertically joined to the equivalent force in the force-curve to estimate the peak force. While measuring the unfolding profile through subsequent force increase scans, we observed two distinct unfolding steps of the talin: one at 9 pN (low force) and the other one at 29 pN (high force) (Fig. 1B). This bimodal mechanical response as an intrinsic functional reporter, revealed that talin could structurally poised to unfold at two different force ranges, suggesting the presence of its two different mechanical conformations.

**Figure 1:**
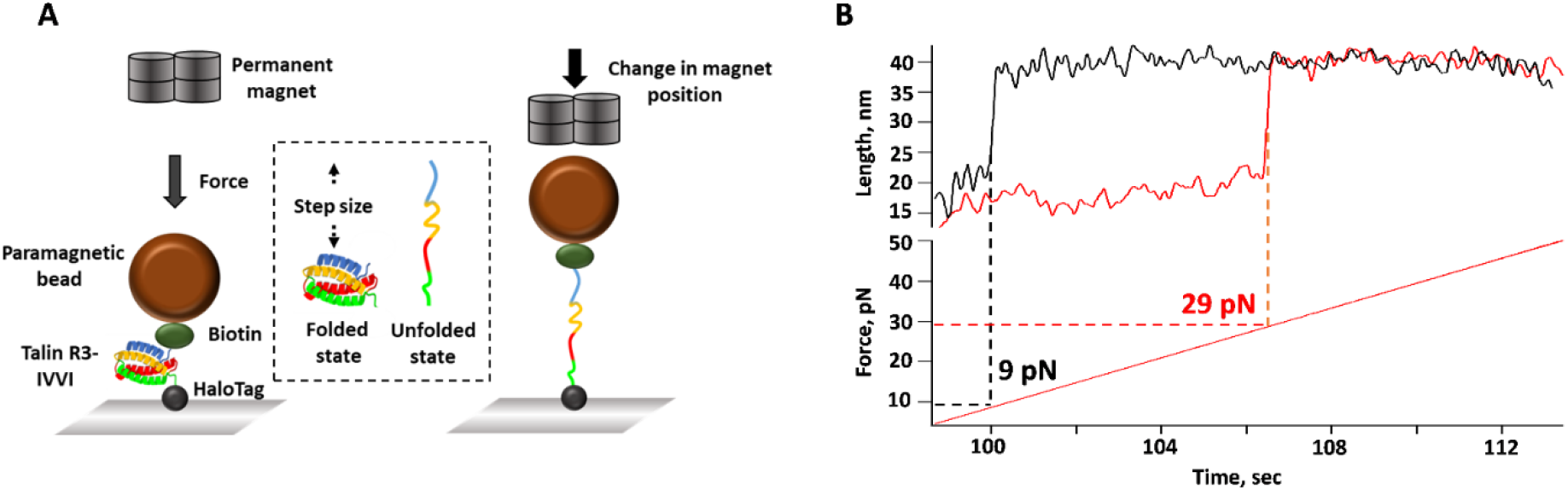
Single-molecule magnetic tweezers experiment: **A. Instrumentation:** The engineered talin construct is tethered between glass surface through N terminal HaloTag and paramagnetic bead with biotin-streptavidin chemistry. Applied force is inversely proportional to the distance between permanent magnet and paramagnetic bead. At high force, talin domain is stretched and the difference between them is defined as step-size. Image is not scaled. **B. Mechanical unfolding of talin domain:** Force at a loading rate of 3.1 pN/s are applied on talin to observe unfolding steps at particular force. Interestingly, talin exhibits a distinct mechanical response by unfolding at two different forces: one at 9 pN force and the other at 29 pN. These different unfolding force range indicates a molecular heterogeneity of talin unfolding under force.

### Mechanical response of talin is modulated by the drug interactions

Given the ability of exhibiting two distinct conformations, we further explored how this bimodal force distribution of talin could be modulated while interacting with the drug molecules under applied tension. Mechanosensitive FA proteins including actin and FAK have been reported as potential targets for therapeutic applications, but how their mechanical responses are perturbed upon drug binding remains elusive. Since talin has been used as a model FA protein in force spectroscopy techniques, we systematically investigated the drug effects on the talin mechanical stability with three drugs: tamoxifen (TAX), letrozole (LTZ), cyanidin 3-O-glucoside (C3G).

To demonstrate the mechanical stability of talin induced by various concentration of drugs, we first studied the mechanical unfolding events of talin without any drugs. Fig. 2A demonstrates the majority of the unfolding force peaks at 8 pN (denoted as low force population) with a very rare events at 32 pN (as high force population). Interestingly, TAX could alter this force distribution in a concentration-dependent manner. We observed talin optimally unfolds at higher force range in the presence of 40 μM TAX (Fig. 2B), above which it starts to decrease to only 19% high force unfolding events at 75 μM TAX (Fig. 2C). By contrast, with cyanidin 3-O-glucoside, we observed a steady increase in the high force population with the concentration (Fig. 3A-3C) and eventually, at 150 μM C3G, all of the unfolding events occurs at high force (Fig. 3D). We have also plotted the high force population against the concentrations for all the three drugs. We observed a steady increase of the high force population with drug concentration, followed by a decrease after the optimal concentration, suggesting a biphasic drug effect on talin mechanical stability. We observed such biphasic effect with both TAX (Fig. 4A, black square) and LTZ (Supplementary Figure 1, black square); however, with C3G a steady increase of high force population was observed (Fig. 4B, black square). This variation in unfolding force population with the drug concentration clearly indicates that drugs affect the mechanical stability by physically interacting with talin. Since drug-modulated stabilization is proposed to occur through stabilizing a specific protein conformation other than its native topology; drug-bound protein even at very high drug concentration, exhibits higher mechanical stability than in its absence. Our single-molecule experiments explored a surprising behaviour of drug-modulated talin mechanical stability, which revealed a generalized question: do drug interactions cause any secondary structure perturbation in talin that might dictate the extent of mechanical unfolding? For a possible explanation, we performed circular-dichroism (CD) spectroscopy to analyze the TAX and C3G effects on the talin secondary structure and observed insignificant change in the secondary structure of the protein (Supplementary Figure 2A). Therefore, this interesting concentration-dependent force modulation might be attributed to a perturbation in the tertiary structure of talin.

**Figure 2:**
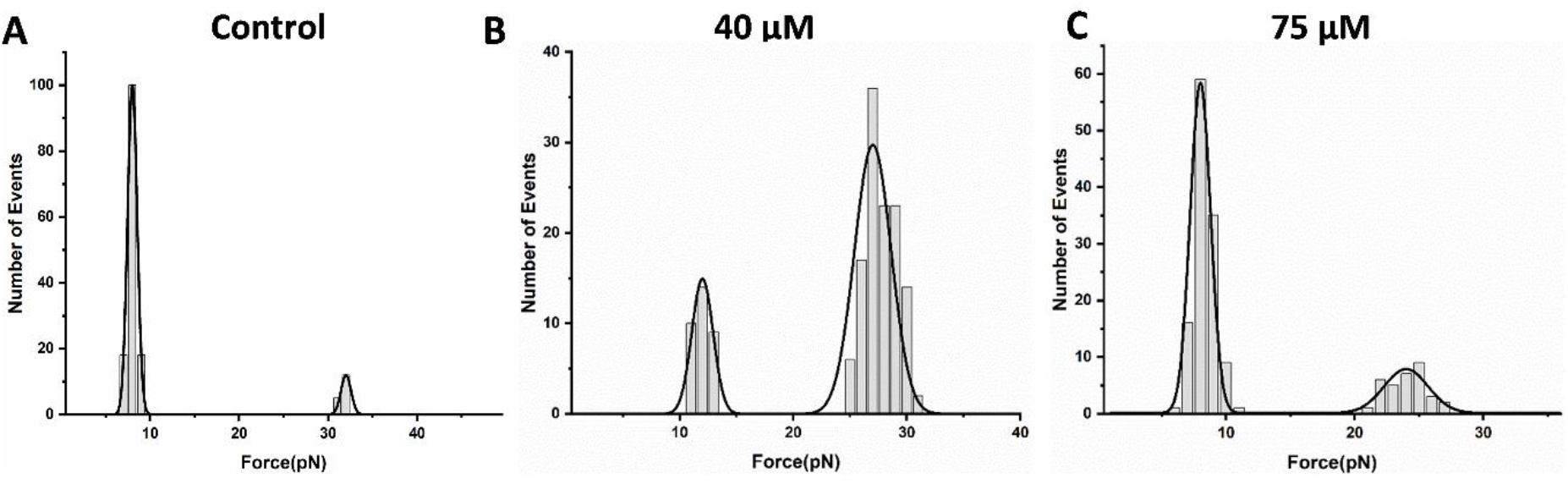
Unfolding force histogram of talin with tamoxifen (TAX): **A.** Unfolding force histogram of talin shows two separate peaks with a very distinct high force population. The solid line is the Gaussian fit to the experimental data. This Bimodal unfolding force distribution has been observed to change upon the addition of TAX. **B.** Interestingly, upto 40 μM tamoxifen, the higher unfolding force population increases with the concentration, **C.** whereas further increasing the concentration to 75 μM again decreases the higher force peak and thus, the talin domain mostly unfolds at lower force.

**Figure 3:**
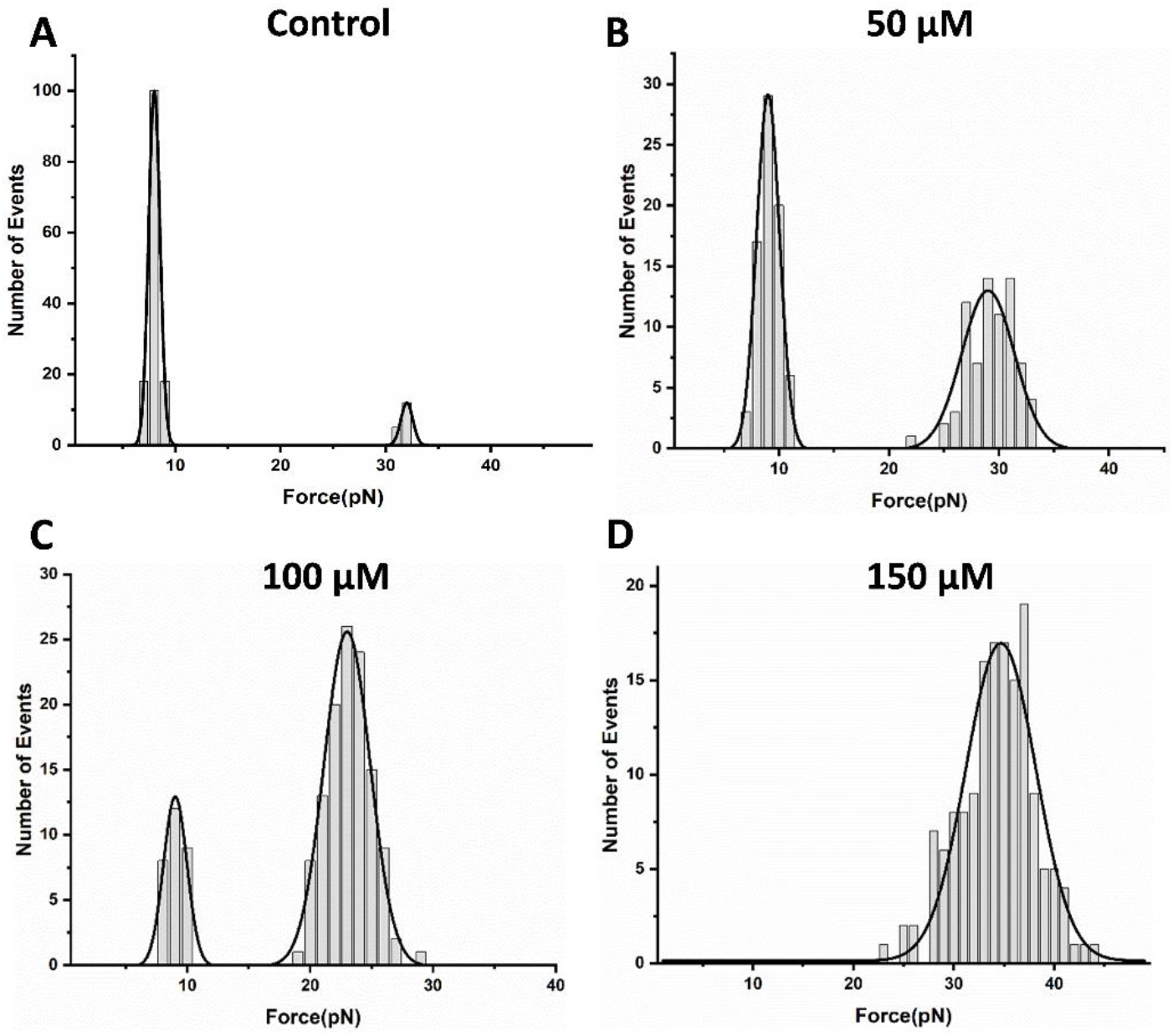
Effect of cyanidin 3-O-glucoside (C3G) on talin domain stability: **A.** Bimodal unfolding force distribution in the control. **B-D.** Relative population of both the low and high force are strongly C3G concentration-dependent. In the presence of 50 μM C3G, most unfolding events occur at low force and fewer of them at higher force. However, at 100 μM, talin unfolding occurs majorly at high force and thus, high force peak becomes larger than that of the small force. And finally, the unfolding events at high force becomes dominant at 150 μM C3G.

**Figure 4:**
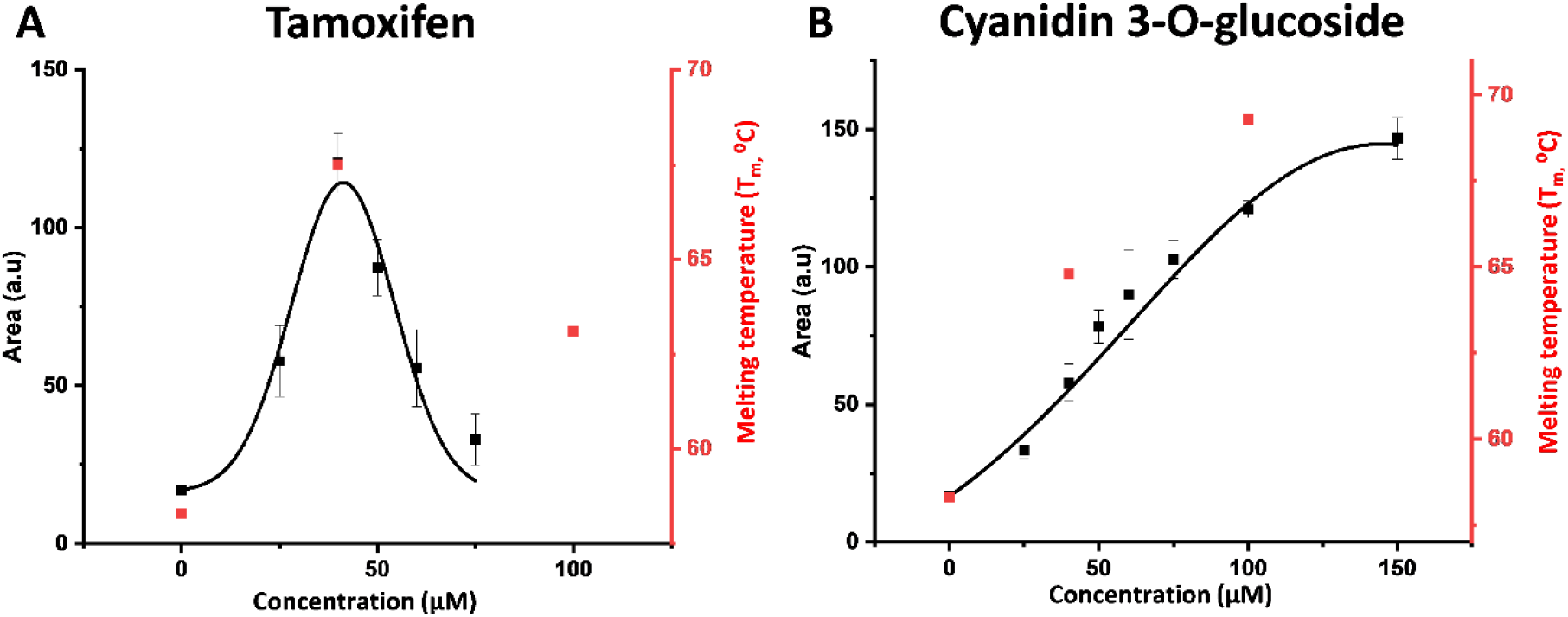
Area analysis and melting temperature: The relative area of high force population (black), obtained from the force distribution curve, are plotted as a function of the drug concentration. **A. TAX:** This area describes that the fraction of mechanically stable domain increases upto 40 μM tamoxifen, followed by subsequent decrease of the fraction with the further concentration increase. Error bars are standard error of mean (s.e.m). Talin melting temperature (red) through its simulated structure, captured at drug-perturbed state follows similar trend with the area curve. Tm of talin has been observed to increase from 58.3° C in native state to 67.5° C in the presence of 40 μM TAX-bound state, however, at 100 μM concentration, it decreases to 63.1° C, following the similar pattern with that obtained with area curve. **B. C3G:** This biphasic effect has not been observed with C3G due to the steady increase in the high force population and interestingly, Tm at different C3G bound state also follow the similar changes as area curve, showing a steady increase in Tm with the C3G concentration.

### Simulations revealed molecular determinant for drug-modulated talin stability

To gain molecular insight into how these drug molecules could affect talin stability, we further carried out force field simulation on GROMACS platform^22^. Since talin mechanical stability varies with drug concentrations, and TAX and C3G display prominent opposite effect; we sought to perform concentration-dependent simulation at 40 and 100 μM concentration of both of these drugs. Direct visualization of MD derived protein-drug interaction as a movie, within simulation timescale provides a range of information (Supplementary Movie 1 to Supplementary Movie 4). Similarly, root-mean square fluctuation (RMSF) indicates the impact of drug interactions on structural stability of proteins through residue flexibility parameter: the region with high RMSF would indicate a more flexible region, while the low RMSF indicates a region with restricted dynamics. Though talin displays minimal structural mobility with C3G, we observed a prominent structural mobility of residues in the presence of 100 μM TAX. From the RMSF analysis, we have observed that regions within 799 and 820 residues are mostly fluctuated in the presence of TAX, followed by perturbation within region within 870 to 880 residues (Supplementary Figure 3). We observed the root mean square deviation (RMSD) of all the systems converged within 50 ns of the simulation time, confirming the stability of the protein-drug interaction (Supplementary Figure 4).

Since protein stability and compactness are interrelated, we measured both the solvent accessible surface area (SASA) and radius of gyration (Rg) to understand the effect of these two drugs. In case of C3G, SASA and Rg has been observed to remain stable with the time at both the concentrations (Supplementary Figure 5A and 6A, green trace). This means C3G do not affect the compactness and stability of talin. However, a markedly increases in both of these parameters has been observed with 100 μM TAX, signifying a loss of compactness in the protein structure (Supplementary Figure 5B and 6B, brown trace). Additionally, influence of 100 μM TAX on protein structure not only increases the surface area but also creates a large cavity (Supplementary Figure 7). To reconcile, we further performed cavity analysis and observed 100 μM TAX creates a large cavity within talin core (Supplementary Movie 2) by largely displacing both helix 1 and helix 3 (H1-H3) and H2-H4 distances from all the three reference vectors (top, center, and bottom) (Fig. 5; Supplementary Figure 8). On the other hand, no subtle change can be observed in any reference position for C3G even at 100 μM. These cavity effects are totally consistent to SASA and Rg analysis. To unravel the basis of this cavity formation, we checked the number of penetrating drug molecules (Fig. 6A and 6B) and surprisingly found that TAX at 100 μM (Supplementary Movie 2) has higher penetration ability than 100 μM C3G (Supplementary Movie 4). At 100 ns time, the number of penetrated TAX molecules is 38, which is zero for C3G (Fig. 6C). Since talin core in hydrophobic in nature, we speculated TAX hydrophobicity could be a factor which became evident from their 6 folds higher logP value than C3G. Even LTZ is almost 2 folds highly hydrophobic than C3G, which could intrinsically link to their strong biphasic effect than C3G, but milder than TAX (Supplementary Table 5). This plausibly indicates a strong contribution of hydrophobicity towards the biphasic effect on talin mechanical response.

**Figure 5:**
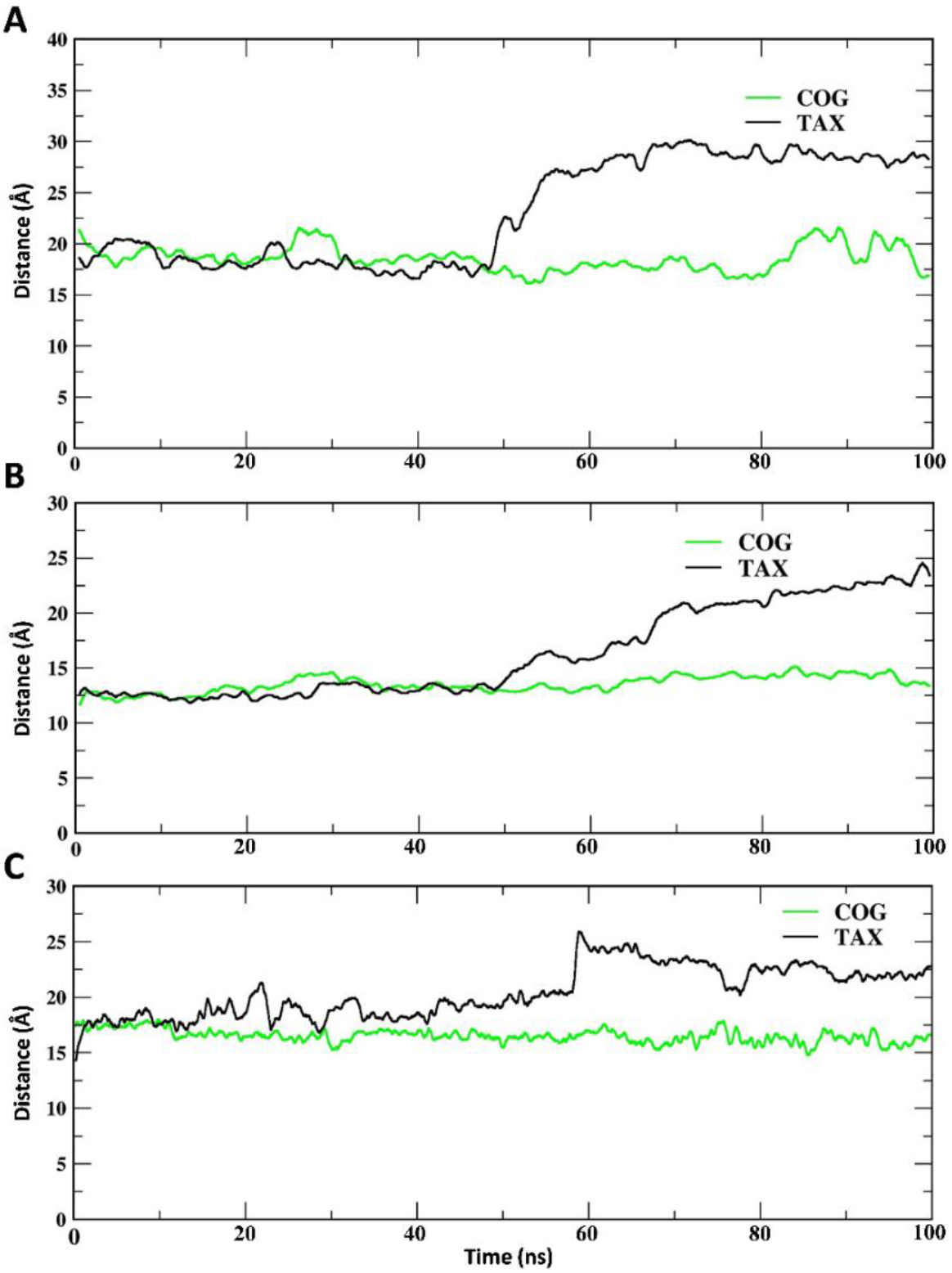
Comparison of helix 1 and helix 3 displacement in the presence of TAX and C3G: Displacement of helix 1 and helix 3 has been observed with both 100 μM TAX and C3G. Reference residues were taken to consider the distance at **A.** top, **B.** center, and **C.** bottom. In the presence of tamoxifen, the helices are displaced and thereby the inter-helix distances are increased with the simulation timescale, signifying a tamoxifen induced mechanical destabilization of talin; whereas with cyanidin 3-O-glucoside, the distances remain unchanged. This indicates cyanidin 3-O-glucoside do not destabilizes talin through separating the helices.

**Figure 6:**
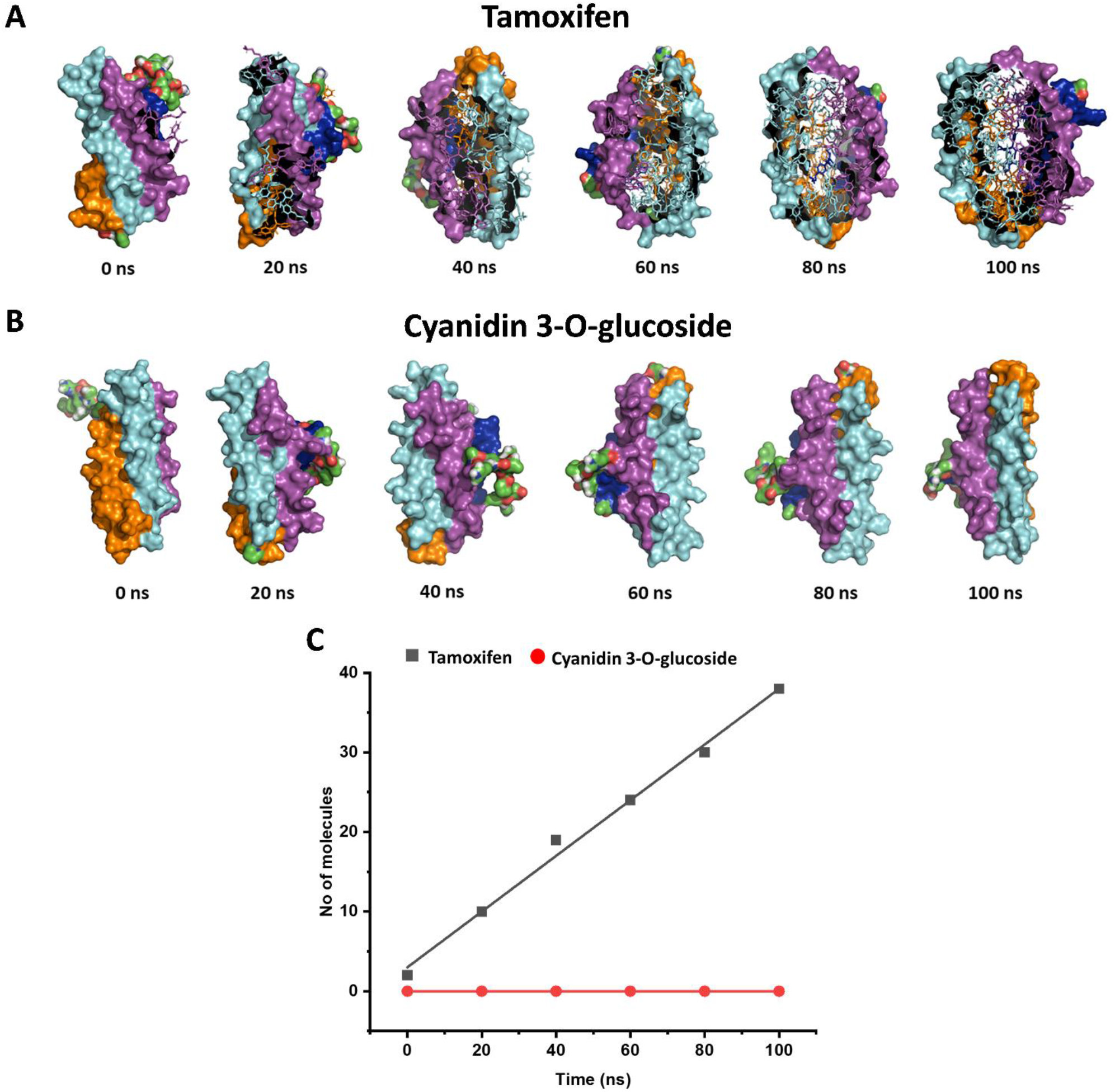
Cavity analysis of talin with tamoxifen and cyanidin 3-O-glucoside: **A. Tamoxifen:** We observed that **A.** TAX increases the cavity within talin protein and penetrates into its helical core, whereas **B.** C3G molecules are not able to penetrate into the talin hydrophobic core and thus, the numbers of penetrating molecules remain unchanged. **C.** Numbers of penetrating molecules for both the drugs are plotted against the simulation timescale.

## Discussion

Our force spectroscopic studies revealed that talin could have bimodal force response while exhibiting the mechanical unfolding; one population unfolds at lower force regime of ~8 pN and other unfolds majorly within ~32 pN. We showed that drug molecules could modulate this bimodal force distribution in a concentration-dependent manner. It is well-known that drugs stabilize the protein in a specific conformation, which could be displayed in local or global conformational change of the protein^23^. We have observed that drug binding to talin, at very high concentration, locks talin in a specific conformation; which display higher mechanical stability than at drug-unbound state. These studied drugs except C3G possess a concentration-dependent biphasic effect on the talin mechanical stability; and interestingly, could be described as a plausible molecular mechanism of biphasic drug effect observed at different cellular processes: metformin displays a biphasic effect on mitochondrial oxygen consumption rate in human induced pluripotent stem cells by modulating the AMP-activated protein kinase signaling^24^. Similarly, studies have revealed that TAX modulates p-glycoprotein mediated intestinal efflux through biphasic dose-response relationship and also danazole-mediated f-actin dynamics^25,26^. Recently, clinical studies have found that LTZ is more effective than TAX against the breast cancer and its recurrence^27,28^. Since talin is significantly involved in the breast cancer malignancy potential through cell migration and invasion^29,30^, our observation on higher talin stability with LTZ than TAX might link to its higher efficiency as a therapeutic. This finding suggests a novel mechanistic perspective on this biphasic effect of drugs on protein stability, especially on force sensitive proteins.

Although previously protein-ligand interaction has been thoroughly studied by force-spectroscopic studies^31–33^, the comparative analysis of drug specificity on the mechanical stability of protein is unknown. Since ligands are well-known to enhance the substrate mechanical stability with the concentration, the observed increase in mechanical stability at first half of the biphasic effect was expected. We experience this trend (for TAX and LTZ) upto optimum concentration; however, further increase in concentration starts to decrease the protein stability. Though talin at very high drug concentration exhibits lower stability than that with optimum concentration; the stability remains still higher than the native talin without drug molecules. Notably, we performed melting curve analysis through its simulated structure generated after each 100 ns simulation along with its native unbound form (Supplementary Figure 9; Supplementary Table 1) and observed both their relative area and Tm change display similar pattern in the presence of both these drugs. For example, Tm of talin is 58.3° C in unbound state, which increases to 67.5° C with 40 μM TAX but eventually declines upto 63.1° C at 100 μM TAX (Fig. 4A), signifying a strong correlation between drug-modulated thermodynamic stability and mechanical stability. However, as a talin-specific drug, C3G exerts a steady increase to both thermal and mechanical stability of talin as a strong indication of binding affinity, which is reconciled from a previous study by Baster et al., where talin has been found to steadily interact with C3G^34^. Notably, these stabilities are attributed to intramolecular atomic contacts and hydrophobic packing of protein^35–38^. Intramolecular contact analysis shows that both the atomic contacts and main chain-main chain hydrogen bonds are reduced in talin with 100 μM TAX, which reduces talin stability (Supplementary Figure 10 and Supplementary Figure 11). Additionally, chord plot analysis shows a loss of interhelix interactions and helix-loop conversion in the presence of 100 μM TAX. However, these changes are not significant with 100 μM C3G, which are indicative of C3G-enhanced talin stability (Supplementary Figure 12). This non-significant changes in secondary structure is also revealed by Ramachandran plot, obtained from simulated talin structure both at unbound and drug-bound conformations (Supplementary Figure 2B-2F). Notably, per residue binding free energy analysis suggests an involvement of Ile, Leu, and Val as major contributors to drug-modulated conformational stability under force (Supplementary Figure 13). Consistent with previous reports that ILV hydrophobic core stabilizes protein conformation^39^, we also found that the total area of hydrophobic core in all the conditions vary accordingly with changing mechanical stability (Supplementary Figure 14, Supplementary Table 2). Mechanically most stable state of TAX-bound talin (talin with 40 μM TMX) shows the lowest binding energy and the least stable state (talin with 100 1μM TMX) shows the highest binding energy indicating the role of hydrophobicity in protein stability. This is also consistent to MM-PBSA analysis with an increased binding energy for lower mechanical stability with 100 μM TAX (Supplementary Table 3). While interacting to the substrate protein, these drug molecules undergoes conformational selection mechanism through preferable binding to force bearing region and therefore, drug-modulated changes in mechanical stability is occurred. From a cellular perspective, it is plausible that drug-modulated mechanical stability might perturb force-dependent interactions of talin with binding partners, either towards the folded state interactors such as Rap1-interacting adaptor molecules (RIAM) and actin or unfolded state interactor vinculin. This could be mirrored in altered migration behaviour of cancer cells through talin-centred mechanosignaling pathways. Finally, we observed that talin-targeting C3G acts in non-biphasic way, stabilizing talin in a consistent way; drugs such as TAX, LTZ which are overall adhesion targeting, not talin, acts through biphasic mode, which could have significant implication in their screening as potential therapeutics.

## Methods

### Protein expression and purification

Talin construct was designed using BamH1, Kpn I, and Bgl II restriction sites in pFN18a expression vector, as described previously ^19,20^. The construct was transformed into Escherichia coli BL21 (DE3) cells, followed by growing them in Luria Broth at 37°C until the O.D.600 reached 0.6. Protein expressions were induced by 1 mM isopropyl β-D-thiogalactopyranoside (IPTG, Sigma-Aldrich) overnight at 25°C. After pelleting the cells, the cells were resuspended in 50 mM sodium phosphate buffer (pH 7.4) supplemented with 300 mM NaCl and 10% glycerol and lysed in the French press (Constant systems). Then the proteins were purified from the lysate with the Ni^2+^-NTA column of ÄKTA Pure (GE healthcare). Then the proteins were biotinylated using Avidity biotinylation kit and recommended protocol, followed by size exclusion chromatography with Superdex-200 increase 10/300 GL column, eluting the protein in Na-P buffer with 150 mM NaCl.

### Flow chamber preparation

The cover glasses (bottom) were sonicated with Hellmanex III (1.5%) and washed with double distilled water. Then it was treated with a mixture containing conc. Hydrochloric acid (HCl) and methanol (CH3OH), followed by conc. sulphuric acid (H2SO4). Then the cover glasses were washed with double distilled water, boiled and dried. Cover glasses were then activated by silanization with an ethanol solution containing 1% v/v (3-Aminopropyl) trimethoxysilane (Sigma Aldrich, 281778) for 15 minutes, followed by several washes with ethanol to remove unreacted silane from the glass surface. Finally, the cover glasses were dried and baked at 65°C. Similarly, coverslips (top) were sonicated with 1.5% Hellmanex III for 15 minutes, followed by washing with ethanol and baked at 65°C for 10 minutes. Then both the top and bottom glasses were sandwiched, separated by parafilm template between the two glasses. After sandwiching, the chambers were vigorously flushed with glutaraldehyde and kept to react for an hour. A PBS solution containing the reference bead (2.5-2.9 μm, Spherotech, AP-25-10) was then passed and incubated for 15 minutes, followed by extensive washing with PBS. Then Halo-Tag (O4) ligand (Promega, P6741) were passed and incubated overnight at room temperature. Finally, to remove any non-specific interaction, chambers were washed with blocking buffer (20 mM Tris-HCl, 150 mM NaCl, 2 mM MgCl2, 0.03% NaN3, 1% BSA, pH 7.4) for ~5 hours at room temperature.

### Single-molecule magnetic tweezers experiments

Single-molecule experiments were performed on custom-made magnetic tweezers built on a Zeiss Axiovert S100 microscope, as described previously^40^. Experiments were performed in a flow chamber, placed on the microscope stage and illuminated with a collimated cold white LED (Thor Labs). The beads were visualized using 63X oil-immersion objective, attached with the nanofocusing piezo actuator (P-725, Physik instrumente). Images were processed through Ximea camera (MQ013MG-ON). Both the piezo positioning and data acquisition were controlled by a multifunctional DAQ (Ni-USB-6289, National Instrument). Forces were applied through the generation of magnetic field by neodymium magnets, which are attached to the linear voice coil (Equipment Solutions) the force can be controlled by changing the position of the magnets. Detail information on force calibration, bead tracking and image processing were discussed previously by Chakraborty et al^41^. Experiments were performed in flow chamber with 1-10 nM protein in 1X PBS buffer (pH 7.2). Drugs were commercially purchased: cyanidin 3-O-glucoside (1151935, Cayman chemicals); letrozole (L6545, Sigma Aldrich); tamoxifen (T5648, Sigma Aldrich). For experiments with drugs, we prepared 250 μM stock solution for each of the drugs in 1X PBS (pH 7.2) and used a range of concentrations. 0-100 μM for LTZ; 0-80 μM of TAX; and 0-150 μM for C3G are used in the single-molecule experiments.

### CD spectroscopy

Far UV CD spectra were carried out using a Jasco J-1500 spectrophotometer. Experiments were performed with 20 μM talin protein in the presence of 40 and 100 μM of both TAX and C3G at far UV spectra, recording over a 200-250 nm range.

### Molecular dynamics simulation

Molecular simulations of protein with TAX and C3G were run using Gromacs 2019.4 package using concentration dependent simulation protocol. Talin structure was taken from PDB ID 2L7A^42^ and IVVI (T809I/T833V/T867V/T901I) mutation was incorporated using PyMOL 2.5.2 version. For the simulation study, GROMOS 54A7 force field was used to study the protein-ligand simulations with SPC/E water model. The protonation states of the talin amino acid residues were determined by H++ server^42^. The systems were solvated with ~32,000 SPC/E water molecules^43^ in a 2 nm cubic box with periodic boundary conditions, while Na+ and Cl-ions were added to reach neutrality and the final concentration of 150 mM. The force fields of the ligands were generated using Automated Topology Builder (ATB) repository. Independent simulation of 50000 steps were carried out for each system. Energy minimization were carried out using the steepest descent algorithm^44^. Initial drug structures were kept randomly in a 10×10×10 nm cubic box. Each complex was equilibrated at 310 K and 1 bar pressure and then was simulated for 100 ns with 2 fs step size. Energy minimization were carried out using the steepest descent algorithm. Each system reached equilibrium at 0.1 ns. Subsequently, the systems were gradually heated up using Berendsen thermostat with a coupling constant of 0.1 ps to reach a temperature of 310 K to perform equilibration in NVT ensemble^45^. Later, the solvent density was maintained using a Parrinello-Rahman barostat (algorithm) with a coupling constant of 0.1 ps at 1 bar and 310 K to perform equilibration in NPT (Number of atoms in the system, pressure of the system and temperature of the system) ensemble, by slowly releasing the restrain on heavy atoms in multiple stages^46^. Lenard-Jones potential was used to calculate the van der Waal’s as well as the electrostatic interactions within the cut-off value of 1.0 nm, and Linear Constraint Solver (LINCS) algorithm was used to constrain all the H-bonds^47,48^. For H-bonding calculations, the cut off length for conventional H-bond was set at 3.5 Å^49^. Other information on simulations has been mentioned in supplementary information. All the results from the simulations were visualized in VMD.

Specific number of drug molecules were kept inside the box to attain the intended molar concentrations (Supplementary Table 4).

### Analysis

All the data acquisition and analysis were performed with Igor Pro 8.0 software (Wavemetrics). Simulation data are analysed with Biovia Discovery studio visualizer. Contacts map and hydrophobic core were analysed by ProteinTools. Thermal stabilities were measured by Scoop program.

## Author contribution

S.C., S.H. design the project. S.C., M.B., D.C. performed the experiments. S.C., D.C., and S.H. wrote the manuscript.

## Acknowledgments

We thank Ashoka University for support and funding. S.H. thanks DBT Ramalingaswami Fellowship and DST SERB Core Research Grant for funding. We thank Dr. Ben Goult (School of Biosciences, University of Kent) for kindly sharing with us the talin clones. We also thank Dr. Abdul S. Ethayathulla (Department of Biophysics, AIIMS Delhi) for helping us with the circular dichroism spectroscopy facility. We are thankful to Dr. Akio Kitao (Tokyo Institute of Technology) for critical discussion about concentration dependent simulation. We also acknowledge BioNome (www.bionome.in)for MD simulations.

## Supplementary information

### Effect of drug binding through molecular dynamic simulation

To check the effect of interactions we performed the RMSD, RMSF, SASA (solvent accessible surface area) and radius of gyration (Rg). The RMSD plots were generated for the protein backbone atoms considering the whole trajectory. While for RMSF, Rg and SASA the simulation window with an appreciable stability index was considered. The data were analysed for the last 50 ns stable trajectory in a window size of 10 ns. The free energy change (ΔG) of the protein was measured using the formula^1^:

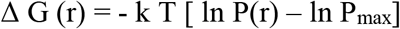

where k is the Boltzmann constant, P is the probability distribution of the molecular system along a coordinate R, and P_max_ denotes its maximum value, which is subtracted to ensure ΔG= 0 for the lowest free energy minimum. We have considered Rg and RMSD for R.

### MM-PBSA analysis

MM-PBSA (Molecular Mechanics Poisson-Boltzmann surface area) approach was utilized to achieve the drug-protein binding free energy calculation^2^. To obtain accurate result, we computed ΔG for last 50 ns in a window size of 10 ns. The free energy of drug binding with the protein was obtained by computing the difference between bound and unbound states employing the following equations^3^:

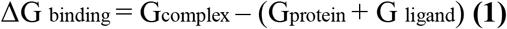

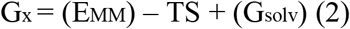

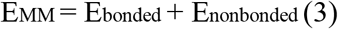

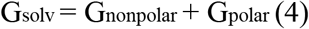

Here, Gcomplex is free energy of protein-ligand complex (PL), (ligand is drug molecules), Gprotein and G ligand are respective free energies of protein (P) and ligand (L). The free energy in any of the bound or unbound state is calculated using the formula (2). Where, *x* represents either PL complex or unbound states namely P or L. The average molecular mechanics energy is calculated using EMM, while TS represents entropic contribution. Finally, *Gsolv* represents the free energy of solvation in binding ligand to protein. The molecular mechanics energy *EMM* was calculated by considering bonded and non-bonded (electrostatic and van der Waal’s) interactions between protein and ligand as shown in the formula (3). While *Gsolv* accounts for linearized Poisson Boltzmann equation for each state (Gpolar) and the non-polar hydrophobic contribution of the system was also considered by calculating for the solvent accessible surface area (4). More complicated entropic contribution was not considered in the analysis with an assumption that the binding of similar compounds with the same protein will have nearly similar entropic contribution, which will be cancelled out during calculations. Finally, the overall binding energy were calculated using:

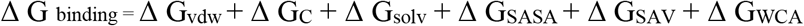

Where, vdw = Van der Waal’s energy, C= coulombic energy, solv = Polar solvation energy, SASA= free energy due to solvent accessible surface area, SAV = energy for solvent accessible volume, WCA = Weeks Chandler Andersen interaction energy.

### Measurement of Protein structural change due to drug binding

Ramachandran plot analysis of talin both at drug unbound and at drug-bound states (as used in the simulation study) were performed. This was done using the Ramachandran plot function of the Biovia Discovery studio visualizer taking the Normal PDB structure of the talin protein and the PDB structures generated from 100 ns simulations in each case. Furthermore, to determine the structural changes of the protein at tertiary level, the chord plots of intramolecular interactions between various secondary structures in the protein with and without the drug molecules were generated using the protein contacts Atlas web program^4^. Additionally, contact map at residue level were generated using ProteinTools^5^ and Biovia Discovery studio separately for overall contacts and intramolecular H-bonding interactions respectively. To check the cavity formation during drug-protein interaction, the varying distance between the helices were measured. We measured the distances between helix 1-helix 3 and helix 2-helix 4 considering the reference top, centre and bottom position of the helices. To measure the number of molecules entering the cavity, snapshots were taken at 0, 20, 40, 60, 80 and 100 ns of the production mdrun. Number of molecules in contact with a particular helix were calculated by taking a cut-off length of 3.5 Å from the helix towards the centre of the protein molecule^6^. Thermal stabilities of the protein in normal and drug-bound condition at different concentration were checked by determining the melting temperature of the Protein using Scoop program^7^. The ILV-hydrophobic core areas for each case were generated using ProteinTools^5^.

**Supplementary Figure 1:**
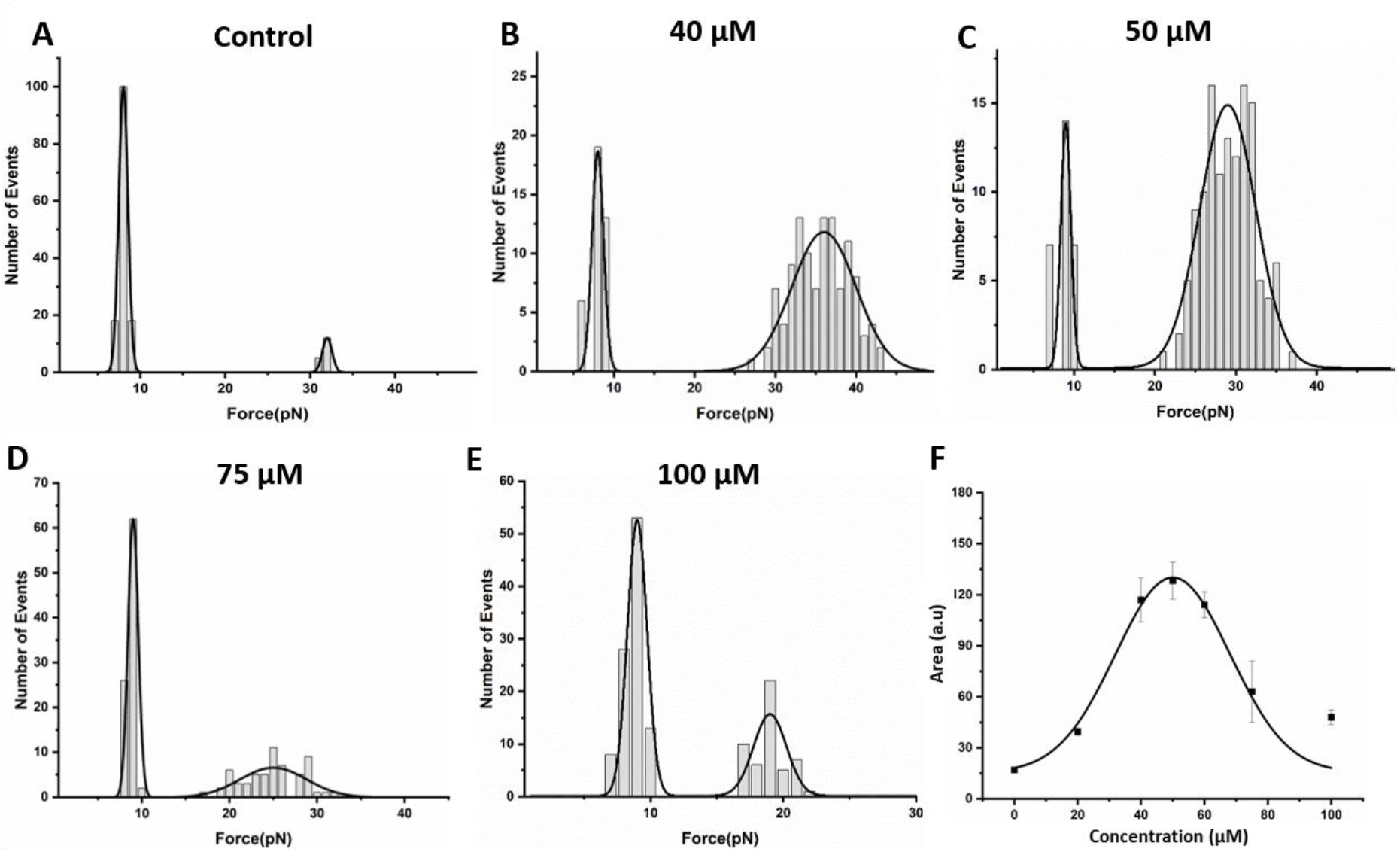
Biphasic effect of letrozole: **A.** Talin shows bimodal force distribution with a rare unfolding event at higher unfolding force. **B-E.** Letrozole exhibit biphasic effect on talin mechanical stability with highest high force population at 50 μM. **F.** The area of talin high force population are plotted as a function of letrozole concentration, showing a biphasic effect on talin mechanical stability.

**Supplementary Figure 2:**
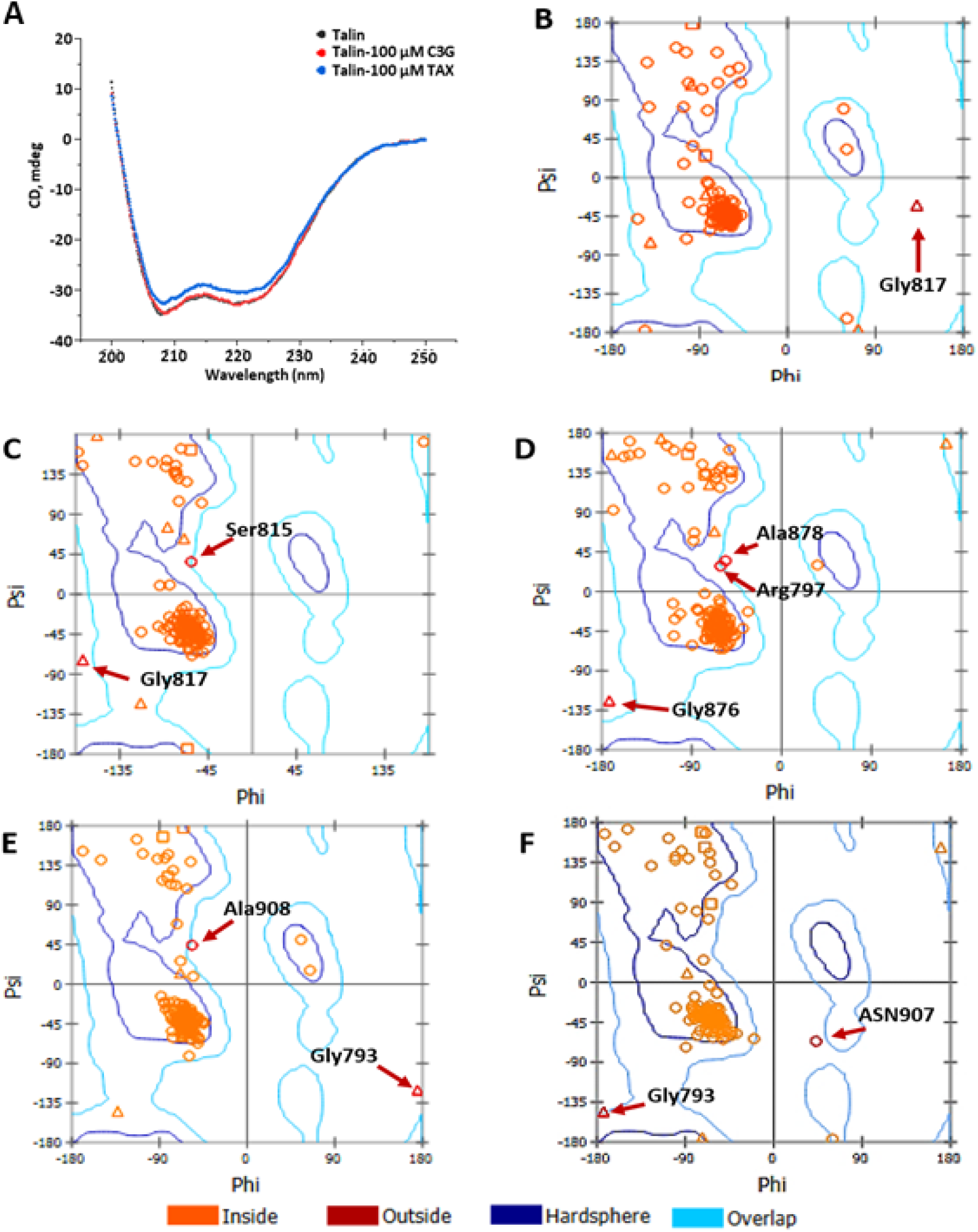
Secondary Structure analysis: **A.** CD spectra of talin (black), talin with 100 μM C3G (red) and 100 μM TAX (blue). **B-F.** Ramachandran plot analysis of the final PDB files generated from each 100 ns simulations of **B.** talin only, and with **C.** 40 μM and **D.** 100 μM TAX. **E.** and **F.** shows the same for talin in presence of 40 and 100 μM C3G, respectively. The inside marks shown in orange represents amino acids in the favoured region of dihedral angles and outside marks in dark red (marked by arrows) denotes the amino acids in disallowed region.

**Supplementary Figure 3:**
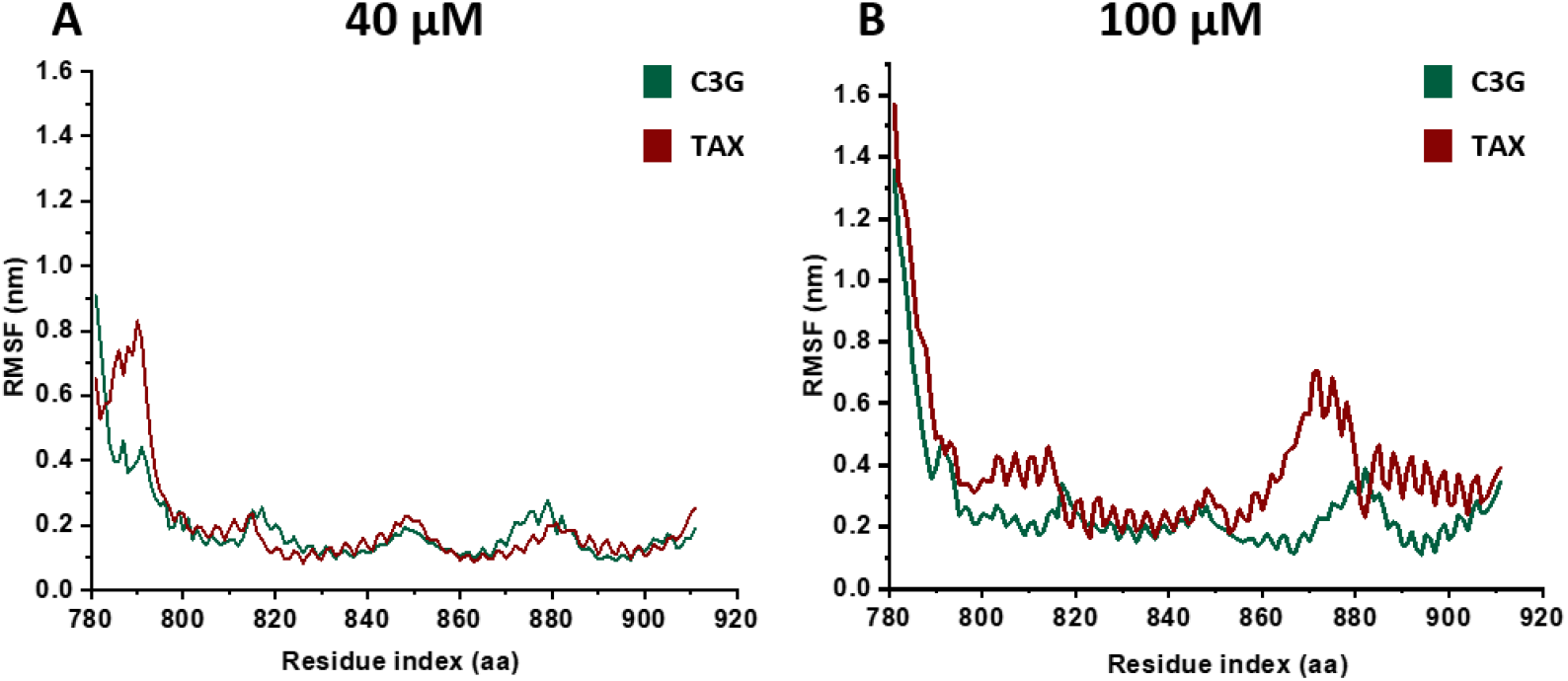
RMSF analysis of talin: **A. 40 μM:** In case of 40 μM TAX (red) and C3G_40 μM (green), the residues in the regions 780-800 (R1), 810-830 (R2), 840-860 (R3), 875-890 (R4) and 905-911 (R5) are mostly fluctuated indicating that either these are the hotspot regions of drug-protein interaction or these regions are more prone to structural change. **B. At 100 μM:** The fluctuation remains almost similar with C3G; but increases significantly at all the region of talin with 100 μM TAX.

**Supplementary Figure 4:**
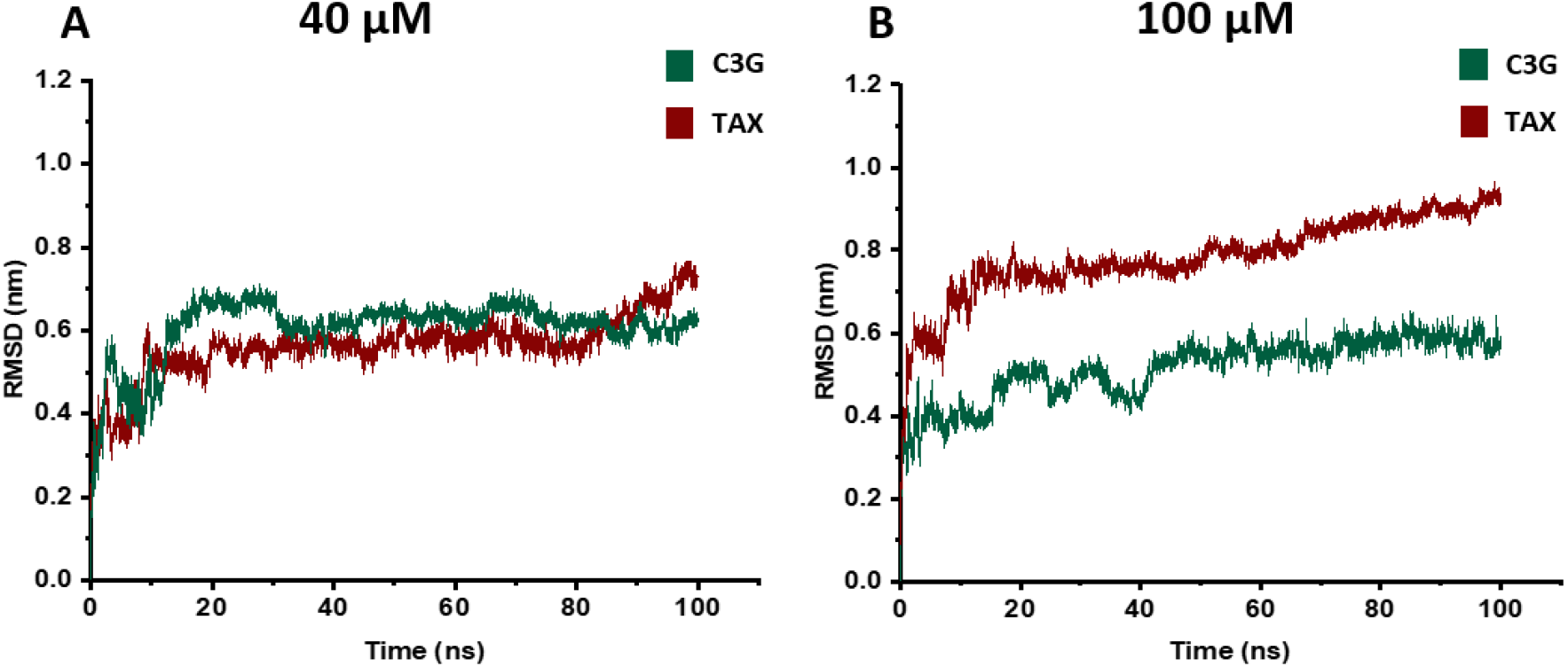
RMSD analysis of talin in the presence of drugs: **A.** The talin-drug interaction is stable with both TAX and C3G at 40 μM concentration, however, B the interaction stability is slightly reduced with TAX at 100 μM concentration.

**Supplementary Figure 5:**
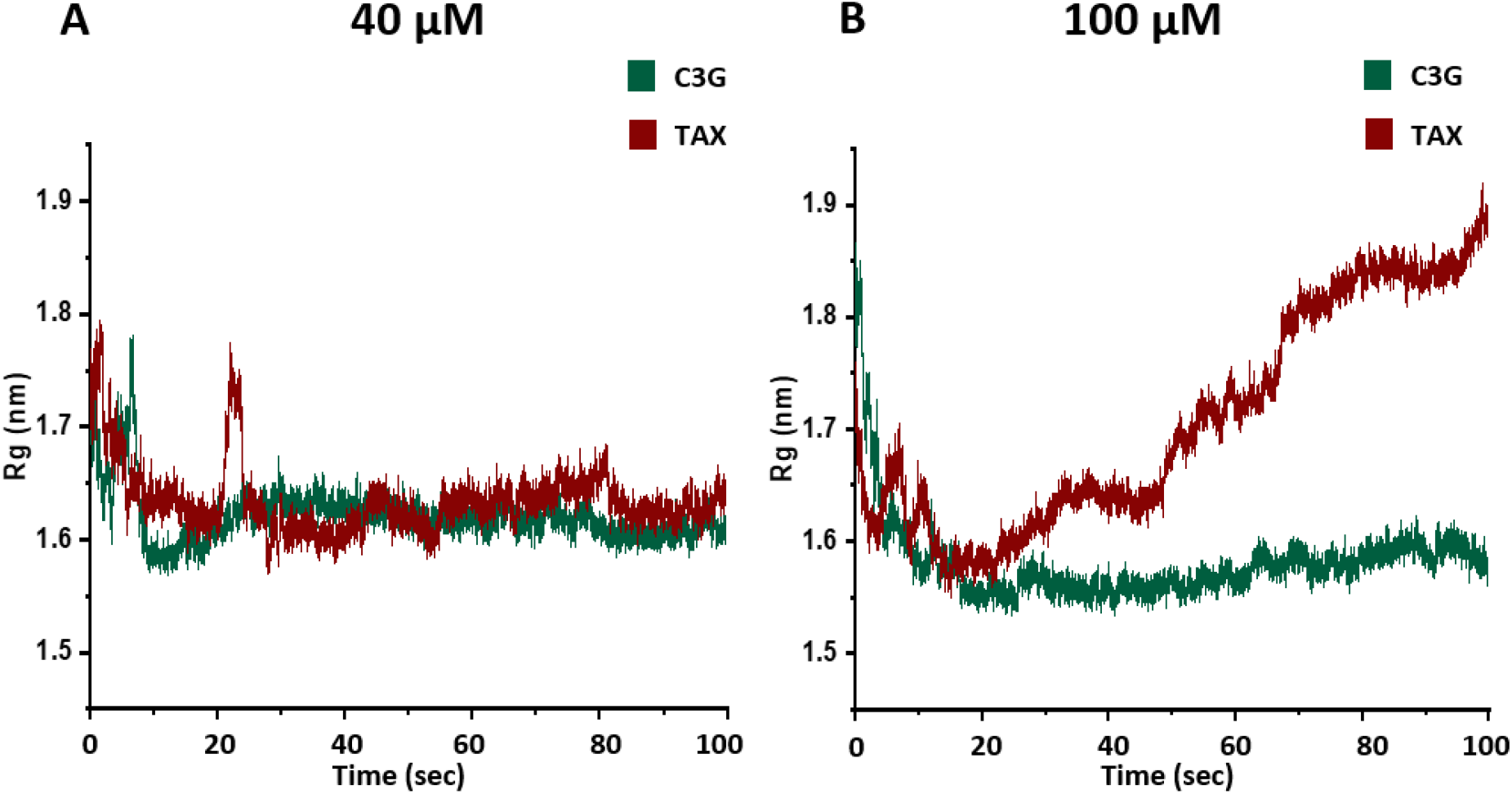
The change in radius of gyration of talin: The change in compactness due to drug interaction is measured through radius of gyration (Rg). **A. 40 μM:** Rg do not change significantly with both TAX and C3G. **B. 100 μM:** At 100 μM C3G, Rg remains stable, however increases ~20% with 100 μM TAX, signifying a loss of compactness in the presence of TAX.

**Supplementary Figure 6:**
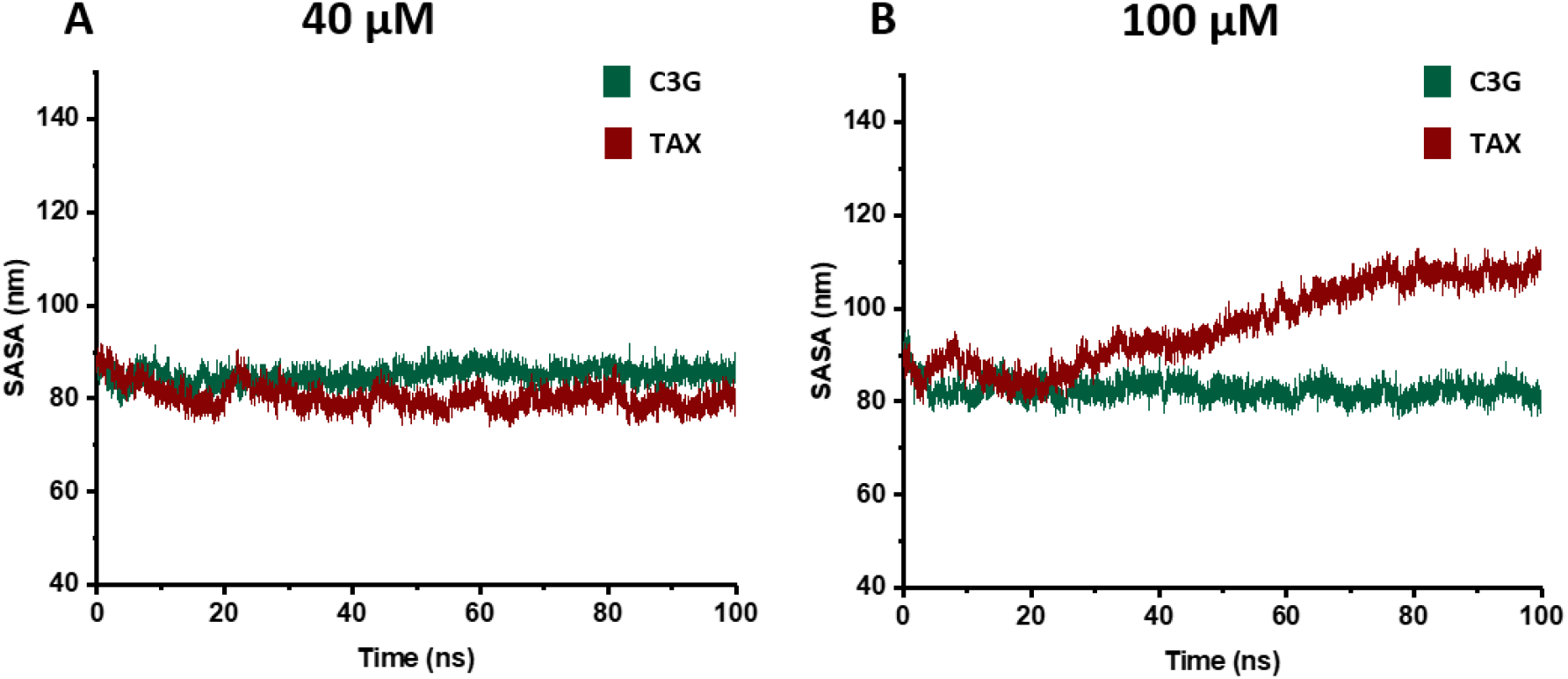
Change in solvent accessible surface area (SASA) of talin: **A.** With both TAX and C3G at 40 μM, no significant change in solvent accessible surface area can be observed. However, **B.** Solvent accessible surface area is significantly increased in the presence of 100 μM TAX, while remained unchanged with 100 μM C3G.

**Supplementary Figure 7:**
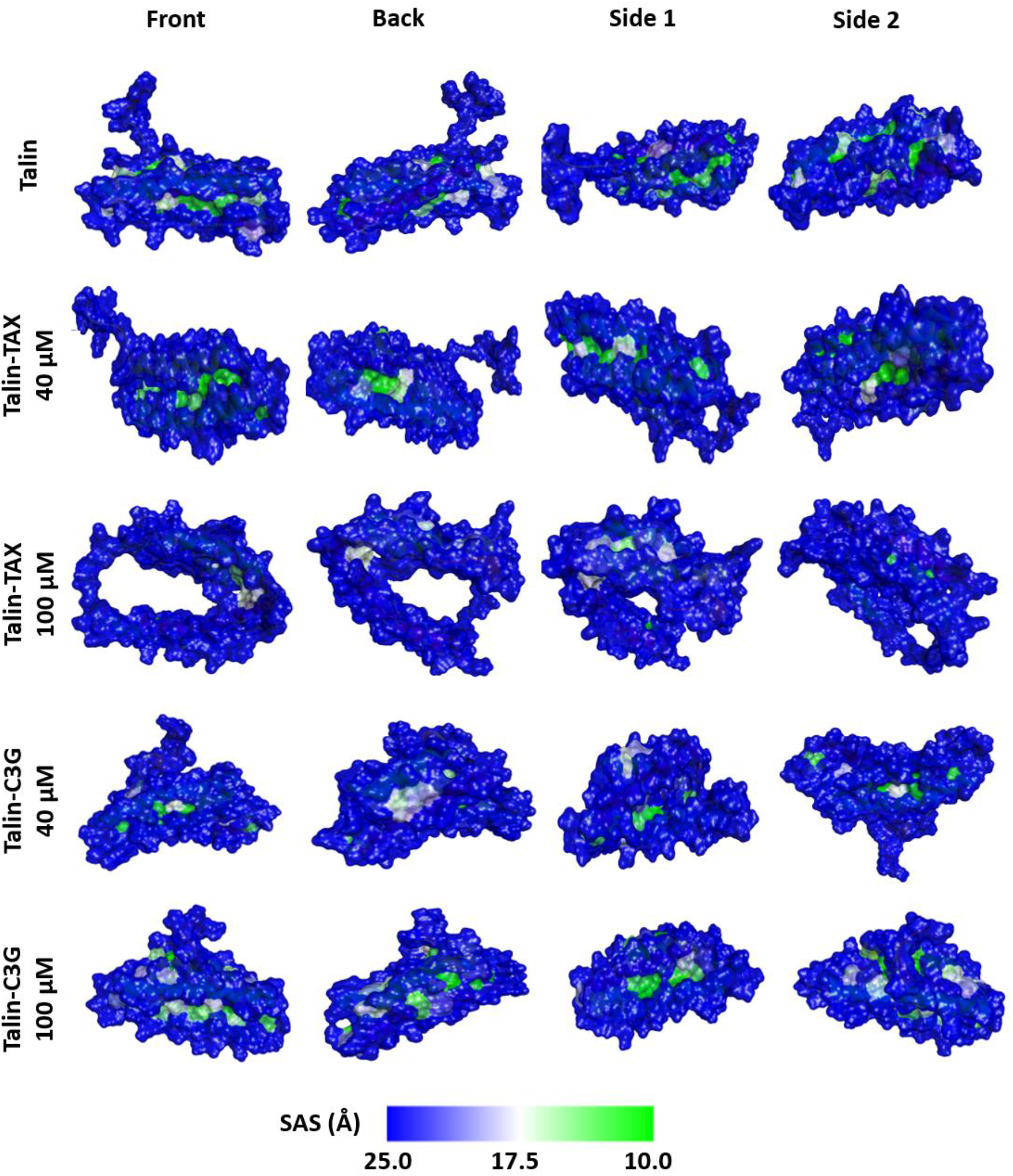
Protein surface model for SASA analysis: Snapshots are taken in different views to show all the solvent accessible and non-accessible surfaces. The amount of solvent non-accessible surface is almost similar at both the concentrations of C3G. Interestingly, it increases significantly while increasing the TAX concentration from 40 to 100 μM.

**Supplementary Figure 8:**
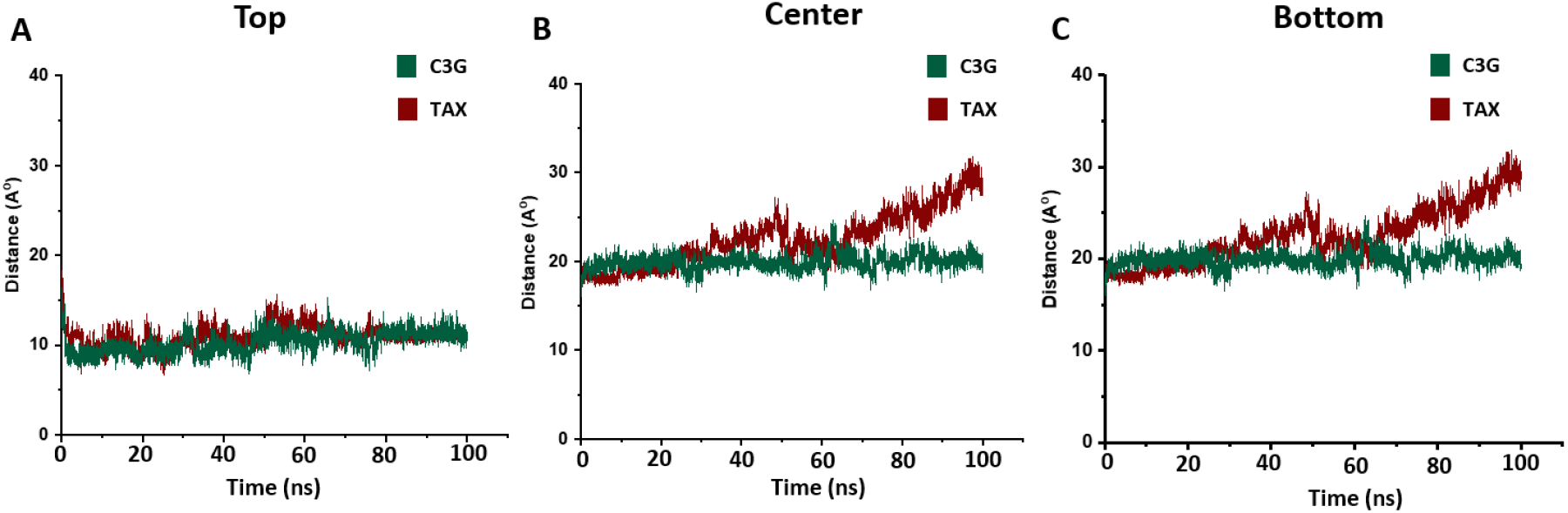
Comparison of helix 3-helix 4 displacement with 100 μM of both TAX and C3G: Reference residues were taken to consider the distance at **A.** top, **B.** centre, and **C.** bottom. No change can be seen in helix distance in top reference position unlike the centre and bottom position, where significant displacements has been observed in presence of 100 μM TAX indicating the role of these positions in cavity formation. The reference residue axes are: 819-884 (top); 832-896 (center); and844-907 (bottom).

**Supplementary Figure 9:**
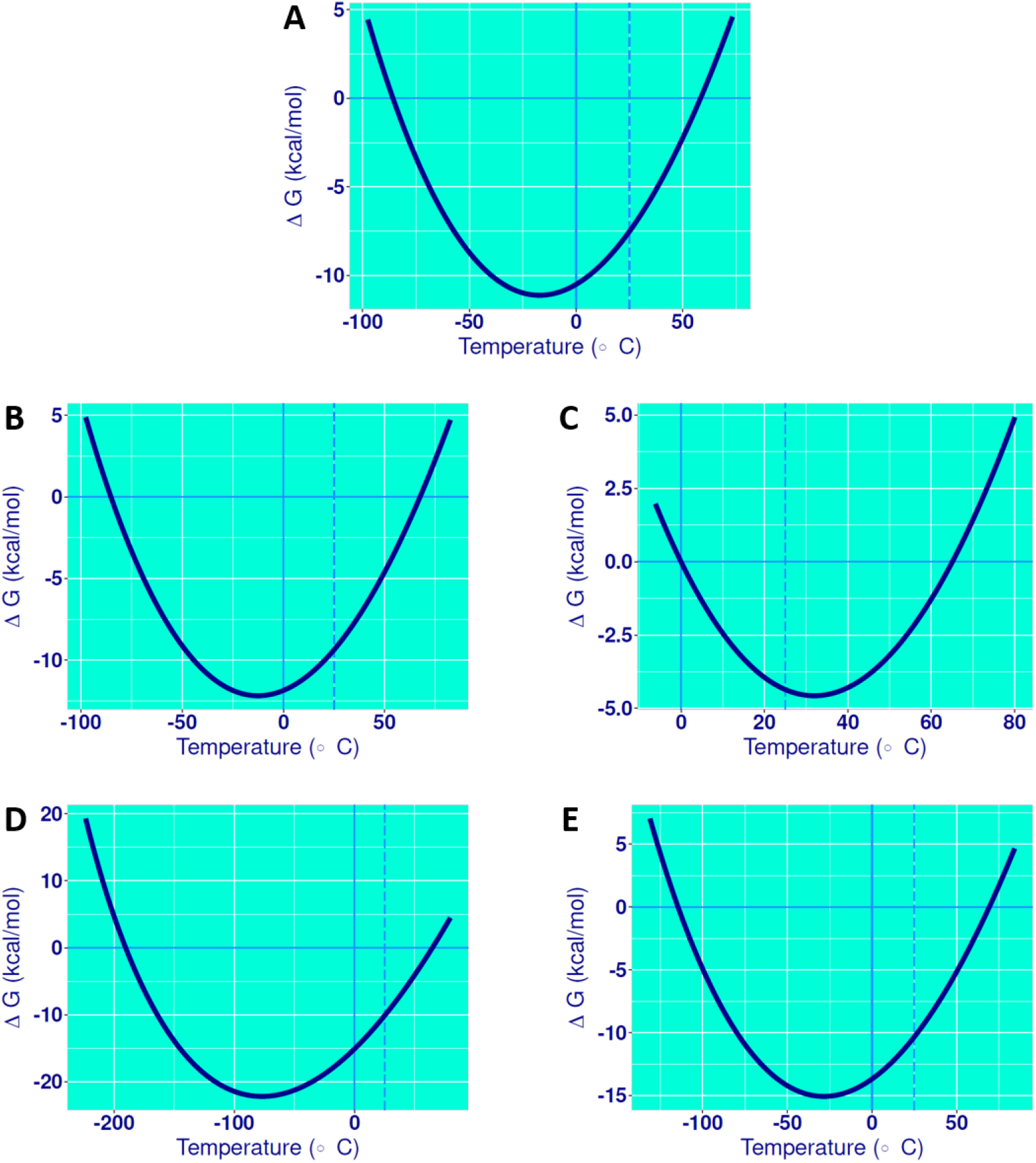
ΔG vs temperature graphs for protein structure talin: (A), in presence of 40 μM TAX (B), 100 μM TAX (C), 40 μM C3G (D) and 100 μM C3G (E). Data from this plot was used to obtain the melting temperature of the protein in drug unbound and bound conditions (Table 3).

**Supplementary Figure 10:**
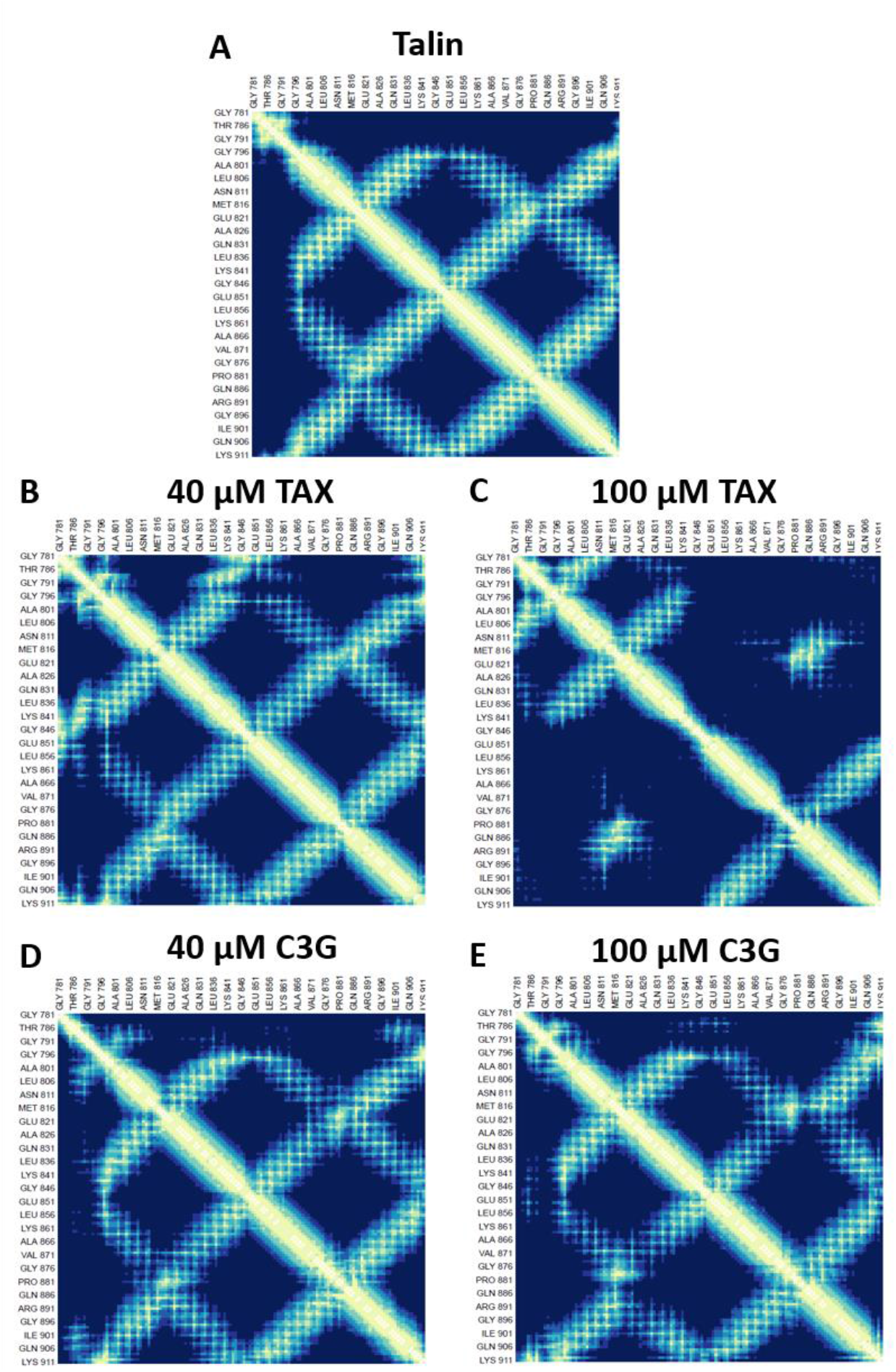
Contact analysis of intramolecular contacts for talin: **A.** Overall contact map for talin shows atomic contacts according to the given scale of contact distances. In case of interaction with **B.** 40 μM TAX and **D.** 40 μM C3G, number of contacts seem to increase. Interactions have decreased significantly in case of (D) TALIN-TAX_100 μM. Further increase in contacts can be seen in case of (E) talin-C3G_100 μM interaction with respect to 40 μM of the same drug.

**Supplementary Figure 11:**
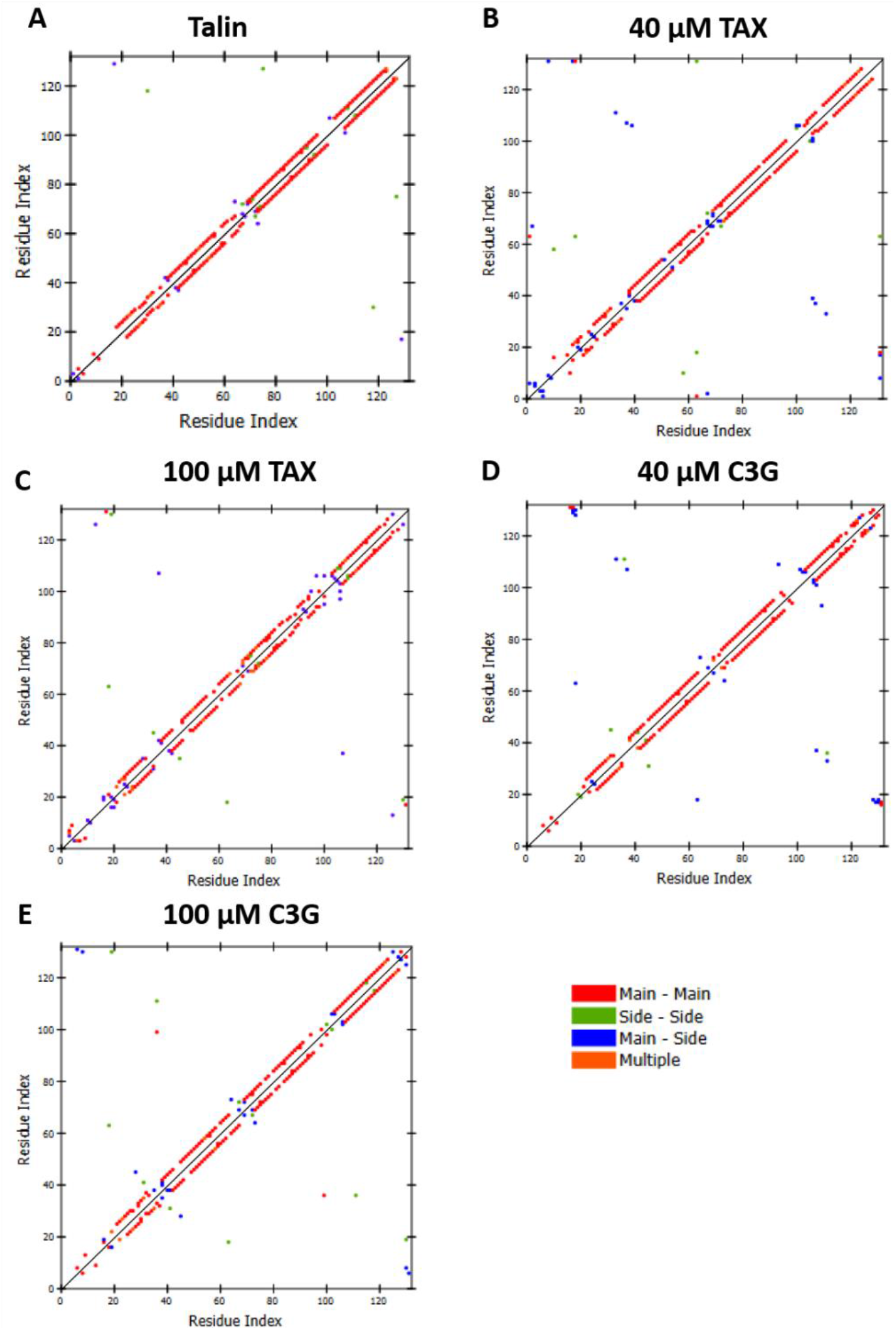
Intramolecular H-bonding contact analysis: Contact plot analysis of various H-bonding interactions among the amino acid residues of the protein at drug unbound state and at drug-bound states at different concentration of TAX and C3G. Amino acid residue numbers are presented as residue index. Interactions are classified as main chain-main chain, side chain-side chain, main chain-side chain and multiple H-bonding interactions. **A.** Overall contact map for talin is nearly similar to that with **E.** 100 μM C3G. Main chain-side chain interactions of talin have increased in case of **B.** 40 μM TAX, **C.** 100 μM TAX, and **D.** at 40 μM C3G. However, talin shows lesser number of main chain-main chain H-bonding interactions with TAX at 100 μM.

**Supplementary Figure 12:**
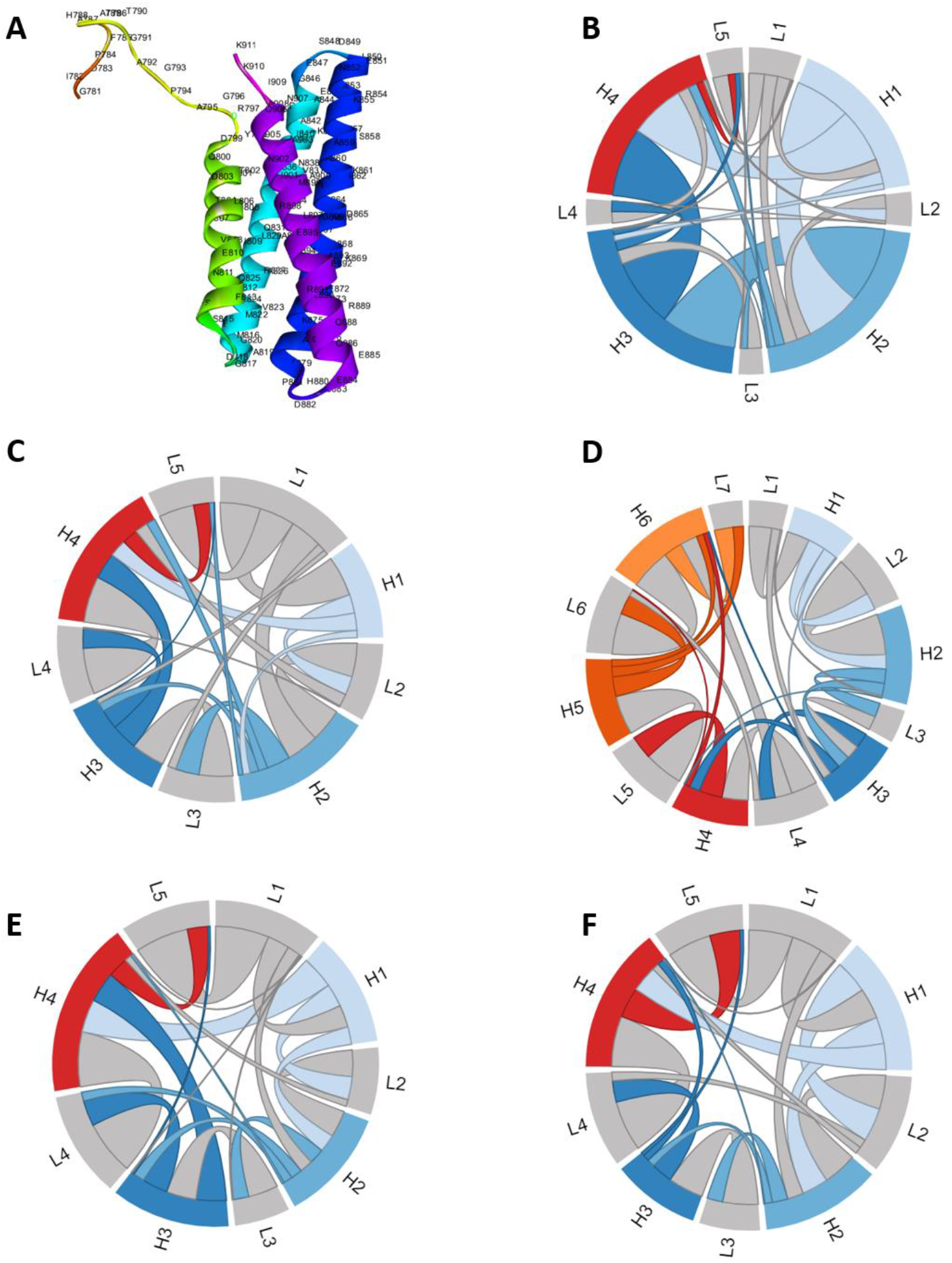
Chord analysis of intramolecular contacts for talin: **A.** The secondary structure of talin labelled with amino acid one letter code. **B.** The chord plots are shown for talin. **C.** Intramolecular interactions maintained in the presence of 40 μM TAX, however, becomes slightly disrupted **D.** with 100 μM TAX, resulting in increase in loop content. **E, F.** Intramolecular interactions with C3G at both 40 and 100 μM concentration. The helix (H) and loop (L) contents are shown below. Different helices are represented in different color whereas grey color is used to mark the loops. **Talin:** L1 = 781-798, H1 = 799-815, L2 = 816-818, H2 = 819-844, L3 =845-849, H3 = 850-879, L4 = 880-883, H4 =884-907, L5 =908-911. **Talin with 40 μM TAX:** L1 = 781-799, H1 = 800-814, L2 = 815-818, H2 = 819-844, L3 =845-849, H3 = 850-879, L4 = 880-884, H4 =885-907, L5 =908-911. **Talin with 100 μM TAX:** L1 = 781-784, H1 = 785-788, L2 = 789-797, H2 = 798-803, L3 =804, H3 = 805-814, L4 = 815-818, H4 =819-836, L5 =837-851, H5 =852-873, L6 =874-882, H6=883-908, L7 =909-911. **Talin with 40μ M C3G:** L1 = 781-802, H1 = 803-814, L2 = 815-818, H2 = 819-846, L3 =847-849, H3 = 850-879, L4 = 880-884, H4 =885-907, L5 =908-911. **Talin with 100 μM C3G:** L1 = 781-801, H1 = 802-814, L2 = 815-818, H2 = 819-846, L3 =847-849, H3 = 850-874, L4 = 875-883, H4 =884-907, L5 =908-911 The numbers are representative of amino acid residues. The strings and chords in the pictures are representative of atomic contacts. The size of the chords varies according to the amount of contact.

**Supplementary Figure 13:**
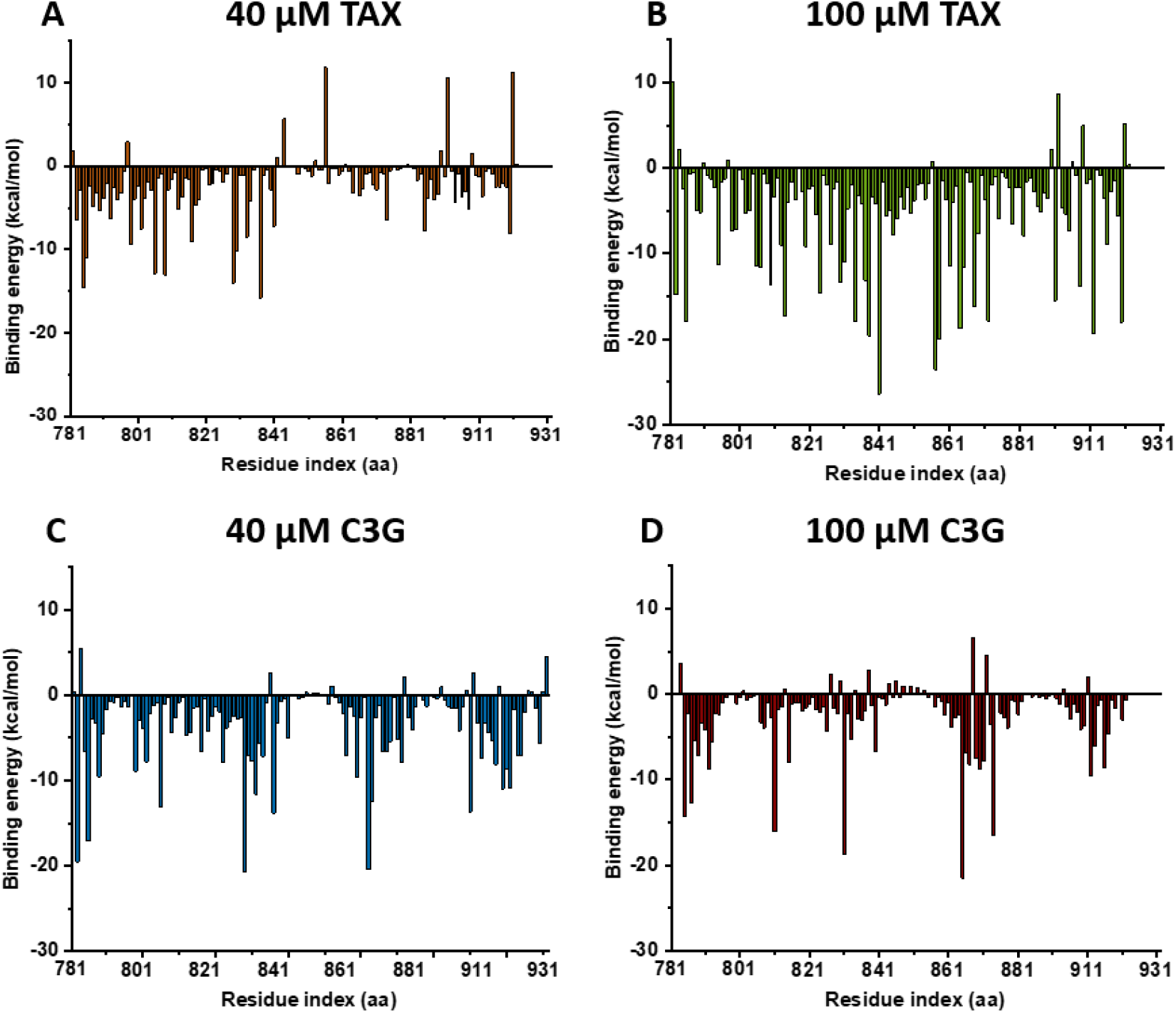
Identification of major contributing amino acids from per residue binding free energy: Per residue binding free energy for amino acids on interacting with **A.** 40 μM TAX, **B.** 100 μM TAX, **C.** 40 μM C3G, and **D.** 100 μM C3G. Fig. C and D shows involvement of almost same amino acids with almost same binding energy. In fig. B, large number of amino acids interact with the drug molecules with binding affinity. This might be due to opening of the protein structure due to interaction with 100 μM TAX.

**Supplementary Figure 14:**
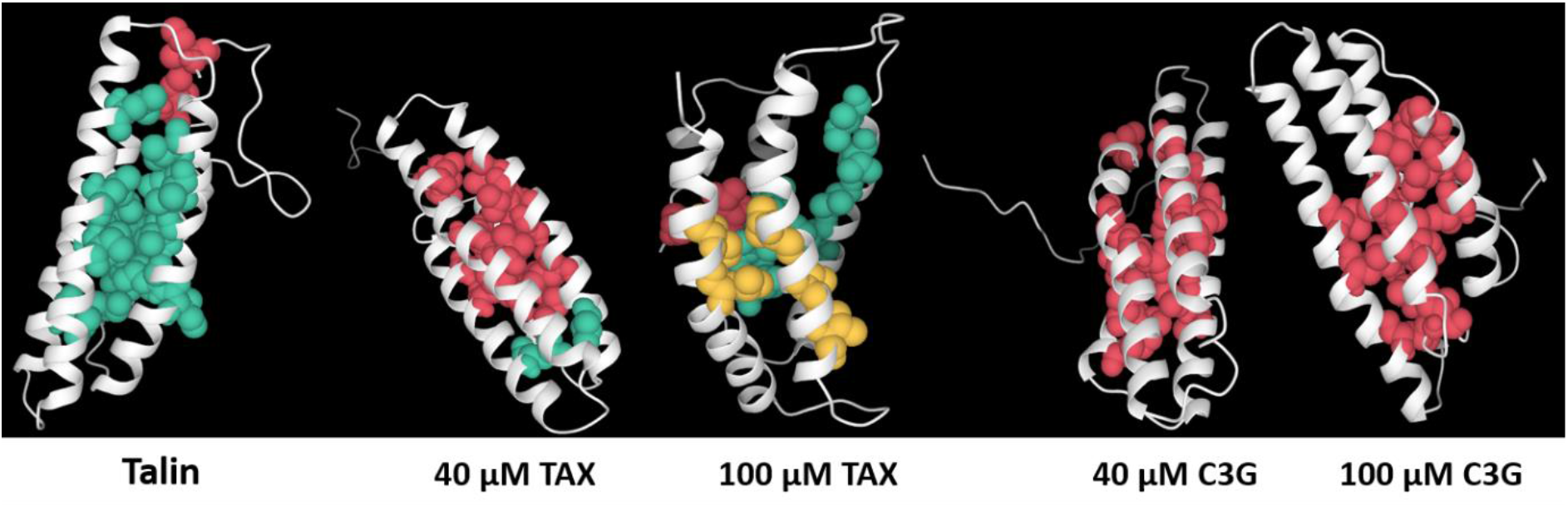
ILV-hydrophobic core formation: Interhelix ILV-hydrophobic core exists in talin. With 40 μM TAX, this interhelix hydrophobic core surface increases, and similarly occurs with 100 μM C3G. However, in presence of 40 μM C3G, the hydrophobic core area remains overall unchanged. But, on interaction with 100 μM C3G, interhelical hydrophobic cores are diminished and strong intra-helical hydrophobic cores are generated. Figures are in accordance with supplementary table 4. The separate cluster of hydrophobic cores are represented by space fill models of different color (red, green and yellow). The protein structure is presented as white secondary structure cartoon.

**Supplementary Table 1:**
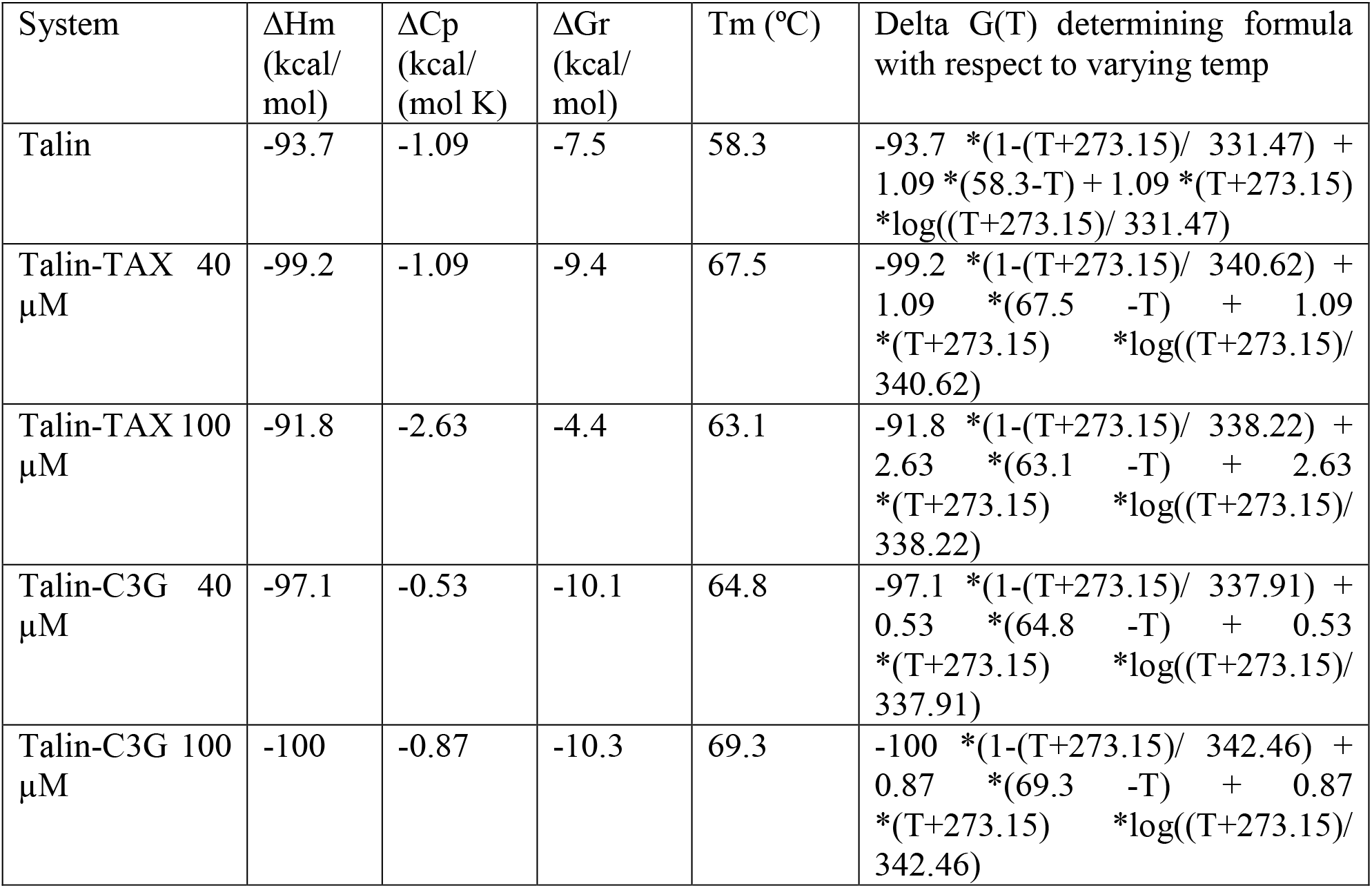
Protein free energy parameters and melting temperature (Tm) in normal and drug-bound conditions at different drug concentration.

**Supplementary Table 2:** ILV-hydrophobic core cluster analysis.

| **Talin** |  |  |
| --- | --- | --- |
| Cluster ID | Area ($\text{\AA}^2$) | Number of contacts |
| 0 | 80.89 | 2 |
| 1 | 1942.14 | 50 |
| **Talin-40 $\mu M$ TAX** |  |  |
| Cluster ID | Area ($\text{\AA}^2$) | Number of contacts |
| 0 | 2221.39 | 48 |
| 1 | 78.46 | 2 |
| **Talin-40 $\mu M$ C3G** |  |  |
| Cluster ID | Area ($\text{\AA}^2$) | Number of contacts |
| 0 | 1966.08 | 54 |
| **Talin-100 $\mu M$ TAX** |  |  |
| Cluster ID | Area ($\text{\AA}^2$) | Number of contacts |
| 0 | 129.65 | 2 |
| 1 | 730.49 | 19 |
| 2 | 355.94 | 9 |
| **Talin-100 $\mu M$ C3G** |  |  |
| Cluster ID | Area ($\text{\AA}^2$) | Number of contacts |
| 0 | 2181.9 | 49 |

**Supplementary Table 3:**
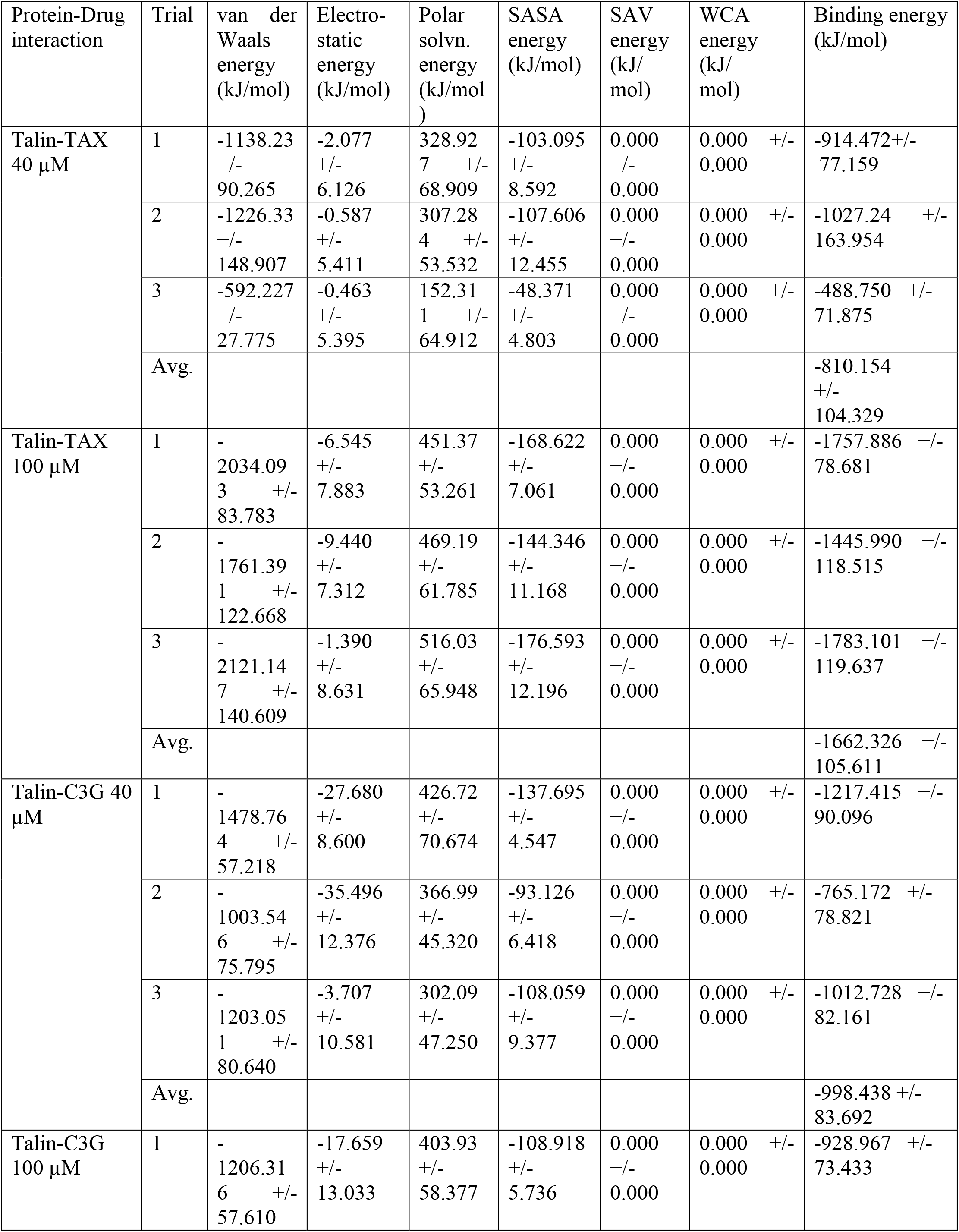
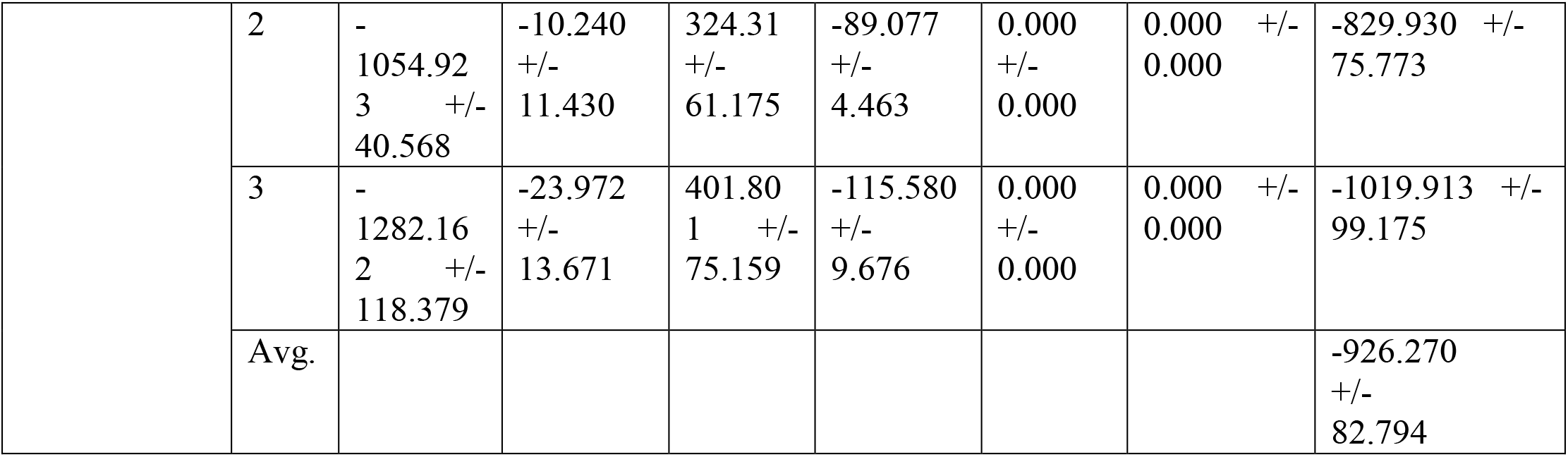
MM-PBSA analysis

**Supplementary Table 4:**
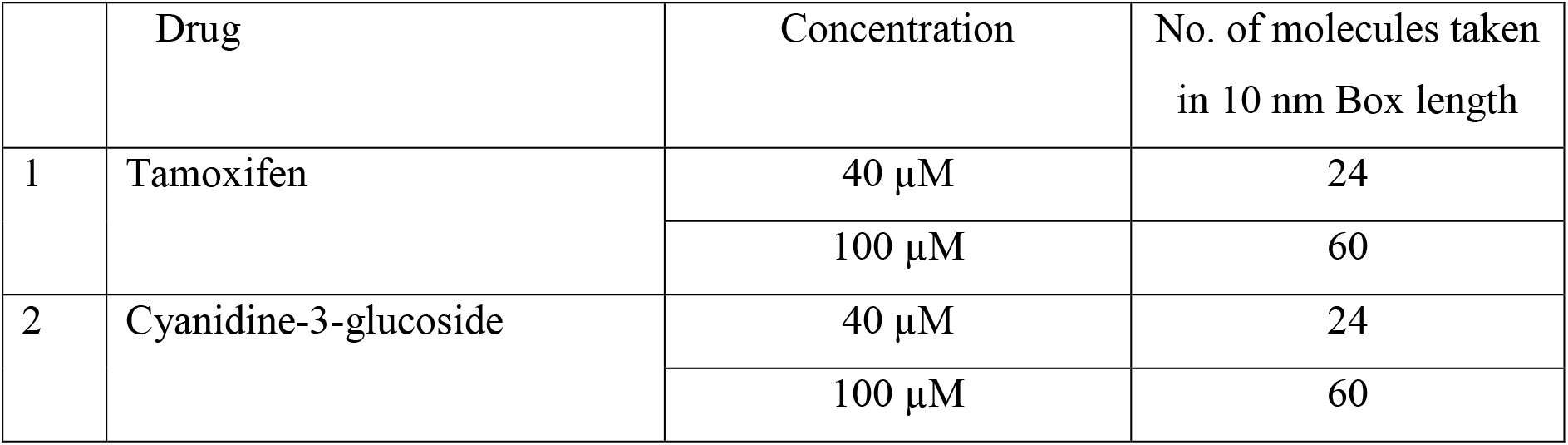
Number of drug molecules for different concentrations.

|  | Drug | Concentration | No. of molecules taken<br>in 10 nm Box length |
| --- | --- | --- | --- |
| 1 | Tamoxifen | 40 μM | 24 |
| | | 100 μM | 60 |
| 2 | Cyanidine-3-glucoside | 40 μM | 24 |
| | | 100 μM | 60 |

**Supplementary Table 5:** logP value of drugs, calculated by ALOGPS 2.1

| Drugs | logP value |
| --- | --- |
| Tamoxifen | 5.93 |
| Letrozole | 1.86 |
| Cyanidin 3-O-glucoside | 0.98 |

**Supplementary Movie 1: Concentration-dependent simulation of talin with 40 μM tamoxifen (TAX):** TAX at 40 μM concentration maintains the inter-helix distance of talin, providing an overall structural integrity to talin. Talin is coloured in green and the TAX molecules are coloured in red.

**Supplementary Movie 2: Concentration-dependent simulation of talin with 100 μM TAX**: 100 μM TAX enhances the cavity formation within talin by displacing helix pairs and penetrates into the helical core. The protein is coloured in green and the C3G molecules are coloured in red.

**Supplementary Movie 3: Concentration-dependent simulation of talin with 40 μM cyanidin 3-O-glucoside (C3G):** C3G at 40 μM concentration maintains the structural stability of talin. The protein is coloured in green and the C3G molecules are coloured in red.

**Supplementary Movie 4: Concentration dependent MD simulation of talin with 100 μM C3G:** Even at 100 μ Mconcentration, C3G maintains the inter-helix distance of talin, providing an overall structural integrity to talin. Talin is coloured in green and the C3G molecules are coloured in red.

